# Herring roe PLs promote SPM biosynthesis in macrophages and a keratinocyte/fibroblast co-culture as model of psoriasis

**DOI:** 10.1101/2025.02.20.639253

**Authors:** Thomas A. Ringheim-Bakka, Amitis Saliani, Tone-Kari K. Østbye, Jennifer Mildenberger, Matthew Dooley, Maftuna Busygina, Mona E. Pedersen, Nina T. Solberg, Jesmond Dalli, Runhild Gammelsaeter

**Affiliations:** Arctic Bioscience AS, Industrivegen 42, 6155 Ørsta, Norway; William Harvey Research Institute, Barts and The London School of Medicine and Dentistry, Queen Mary University of London, Charterhouse Square, London. UK. EC1M 6BQ; Nofima AS, Osloveien 1, 1433 Ås, Norway; Møreforsking AS, Borgundveien 340, 6009 Ålesund, Norway

**Keywords:** drug therapy, inflammation, skin, phospholipids, fish oil, lipid mediator, marine phospholipid esters, herring roe oil, psoriasis

## Abstract

Psoriasis is a chronic immune-mediated inflammatory disease (IMID) affecting the skin presenting with both local and systemic inflammation as part of its pathophysiology. An oil rich in phospholipids extracted from herring roe has been shown to have immunomodulatory properties and to improve clinical symptoms and impact inflammatory cytokine pathways in psoriasis in a clinical trial. The lipidic nature of herring roe oil (HRO) and its high content of polyunsaturated fatty acids suggests involvement of lipid mediator pathways for the observed alleviation of psoriatic inflammation. Of particular interest is the super-family of lipid mediators termed specialized pro-resolving mediators (SPMs), due to their involvement in resolution of inflammation and subsequent return to homeostasis. We therefore explored the influence of HRO and its phospholipids on lipid mediator and SPM biosynthesis in IFN-γ and LPS-stimulated human monocyte-derived macrophages and an IL-17A-stimulated keratinocyte/fibroblast co-culture. Lipid mediators including SPMs were quantified from cell supernatants using a validated LC-MS/MS protocol. In these experiments we observed broad SPM biosynthesis with dominant upregulation of RvE2 and RvE3 in both cell systems and upregulation of DHA-derived SPMs such as RvD2 and PDX. Observations of PCTR2 in macrophage cell supernatants also indicate activation of reparative pathways upon treatment with HRO. In conclusion, we observed promotion of SPM biosynthesis associated with a shift towards a protective and possibly reparative macrophage phenotype as well as promotion of biosynthesis of pro-resolving lipid mediators in a skin cell co-culture, thus demonstrating a possible mechanism for resolution of inflammation in the skin niche using HRO.

## Introduction

Inflammation is a physiological process that has been described by scholars for centuries, even millennia, and inflammation was famously described two thousand years ago by Aulus Cornelius Celsus (25 BC – 50 AD) through the four cardinal signs of redness, swelling, heat, and pain (1–3). With the progress of science, including growing access to modern powerful analytical instruments, an enormous amount of work has since then been put into expanding the knowledge of initiation and perpetuation of inflammation by describing the actions of cytokines and chemokines, as well as the pro-inflammatory eicosanoids (e.g. prostaglandins and leukotrienes) and their importance in driving inflammatory processes (4–6). It has only been in the most recent decades that scientists have been able to elucidate the active processes such as the lipid mediator class switch that are responsible for containing inflammation and restoring tissue homeostasis (4,7). Where before it was thought that resolution of inflammation was a passive process that took place through gradual dilution of chemoattractants, it has now been shown that these are active processes regulated through the expression of a superfamily of lipid mediators termed specialized pro-resolving mediators (SPMs) (2,5,6,8). SPMs are potent and transient small molecule signaling agents that are endogenously biosynthesized from polyunsaturated fatty acids (PUFAs) and that, as implied by their name, promote the shift towards resolution of inflammation (4,6). The biosynthesis of SPMs from their parent fatty acids (FAs) is regulated by cell type and phenotype, enzyme expression, and substrate availability (2,9–11). SPM signaling and regulation result in, amongst other, the reduction of polymorphonuclear leukocyte (PMN) infiltration (6), the promotion of efferocytosis/phagocytosis (6), and promotion of tissue regeneration (12,13).

The lack of, or incomplete, resolution of inflammation perpetuates what should have been appropriate and self-limiting inflammatory processes and has been linked to chronic inflammation and multiple disease states (14). A group of chronic inflammatory disorders with unknown etiology that share common inflammatory pathways has been collectively termed immune-mediated inflammatory diseases (IMIDs) (15). IMIDs are a major public health issue with a global incidence of over 67 million cases (16). IMIDs can affect a number of organs and tissues, including the skin, and can have a profound impact on quality of life, and often present with comorbidities (17–19). In 2019, 6.84% of new IMID cases were psoriasis (16), a chronic inflammatory skin disease which presents with both local and systemic inflammation as part of its pathophysiology. Hippocrates (460 – 377 BC) described the symptoms of psoriasis as *psora* (itch), and *lopoi (*scaling) and recommended treatment with tar followed by sun exposure (20). It is now known that although IMIDs display various symptoms depending on the organ they are affecting, IMIDs share an underlying pathogenesis and, indeed, many of the same drugs are therefore used to treat several of these diseases (21,22). Insights into the molecular basis of inflammatory disease has led to therapies targeting specific IMID-associated immune and inflammatory pathways, with recent focus on developing drugs targeting the cytokines IL-17 and IL-23 (23,24). However, due to their intricate involvement in cessation of inflammation responses, imbalances in lipid-mediated regulation of resolution/inflammation due to dysregulated lipid mediator and SPM biosynthesis and signaling have also been proposed to play a role in the pathophysiology of IMIDs (25–27), and it has been shown that intervention with PUFAs (28) or SPMs (29,30) may hold promise in correcting these imbalances. Lack of immunoresolution due to dysregulated SPM biosynthesis may arise both from dysregulated enzyme expression as a consequence of the disease state (31) or from substrate competition from pro-inflammatory precursors released from membrane phospholipids (PLs) (32).

Marine PUFA oils have traditionally been considered to have anti-inflammatory properties and have been reported to have potential in treatment of selected IMIDs (33,34). Furthermore, supplementation with marine oils has also been reported to promote SPM and SPM precursor biosynthesis (35–37), where biosynthesis and downstream SPM profiles changes according to different marine oils (37). The difference in conversion between oils implies that the pharmacology may be impacted by other parameters, such as lipid composition and oil source, rather than just the PUFA content/ratios. One marine oil of interest in this regard is herring (*Clupea harengus*) roe oil (HRO), where the nature of the raw material used for extraction causes HRO to have a high content of membrane phospholipid esters enriched with marine PUFAs (PEHeRo = phospholipid esters from herring roe). The benefits of marine PLs have been described previously in literature as having anti-inflammatory effects (38–40) and reports also imply that marine PLs could be superior to other marine lipid forms with regards to bioavailability and active uptake across the blood-brain barrier (41–45), neuroprotective properties (46), and their ability to reduce lesion progress in atherosclerosis (47).

The long-chain PUFAs (LC-PUFAs) provided by PEHeRo in HRO are primarily docosahexaenoic acid (DHA) and eicosapentaenoic acid (EPA) in a ratio of approximately 3:1. As reported for other marine PLs, the fatty acids in PEHeRo have also been shown to be readily bioavailable and supplementation has shown a reduction of plasma and RBC PC n-6/n-3 ratios (48), as well as a higher acute uptake of DHA and EPA when compared to a triacylglycerol source in a head to head study (49). The bioavailability of lipids and their corresponding fatty acids upon administration is relevant for lipid mediator biosynthesis, as LC-PUFA supplementation has been shown to increase blood and plasma levels of lipid mediators (50–53). Being able to increase blood levels of lipid mediators through supplementation coincides with observations of positive effects on inflammation through direct supplementation of lipid mediators (31,54,55).

In addition to the general reports on benefits of marine PLs pertaining to inflammation, HRO specifically has been shown to display effects on peripheral inflammation (56) and has been shown to dampen psoriasis *in vivo* in a clinical trial (57–59). The dampening effect of HRO on psoriasis has been shown in human trials where oral daily treatment with HRO gave a statistically significant improvement of mean Psoriasis Area Severity Index (PASI) score (57,58). *Post hoc* analyses of blood samples from the study by Tveit *et al.* (57) further support the beneficial effects of HRO for the dampening of psoriasis by showing that HRO had a beneficial effect on immune cell activation and influenced the immune cell profile by, amongst others, reducing CCL2 over time and leading to an increase in IFN-γR1 (59). The *in vitro* experiments performed by Mildenberger *et al.* (60) further corroborate the potential benefit of HRO in the treatment of psoriasis by showing that HRO limits secretion of IL-23 in macrophages in a dose-dependent manner and limits secretion of IL-17 from CD^4+^ T-cells. Thus, this shows that HRO positively impacts immune cells and skin cells impacted by psoriasis in multiple ways.

To understand how HRO exerts anti-inflammatory actions, we have studied its impact on lipid mediator biosynthesis in several cell systems. We have studied the effects of HRO on SPM biosynthesis in primary human monocyte-derived-macrophages (MDMs), as these cells have a role in all inflammatory processes and are known to readily express SPMs as part of their signaling in inflammation regulation. Macrophages are also known to change their phenotype towards more anti-inflammatory phenotypes upon stimulation with lipid mediators (61), which is followed by a shift in lipid mediator biosynthesis. In addition, the effects of HRO on SPM biosynthesis in a skin cell model consisting of a co-culture of IL-17A-treated keratinocytes and fibroblasts, which mimics psoriasis, were studied (62). Current knowledge about SPM biosynthesis in skin cells is limited. Therefore, the study aimed to elucidate the role of these cells in SPM and lipid mediator biosynthesis during resolution of inflammation. Studying both the immune cells and skin cells simultaneously may give an image of the cell-cell interplay between these cell types locally in the skin niche. Lastly, we also mapped the lipid mediator content of HRO and isolated PEHeRo to have a greater understanding of which mediators were presented to the cells exogenously compared to the output after treatment, and if any of the identified lipid mediators has described roles in specific interleukin pathways relevant to the pathogenesis of inflammatory diseases including psoriasis.

## Materials and methods

### Preparation of HRO and PEHeRo

All lipids were stored under inert gas blanketing (N_2_) at any step. The nitrogen gas blanketing was only suspended during transfer of materials and during autoflash purification.

To produce HRO, immature herring roe (IHR) was dried under reduced pressure (until H_2_O <20% w/w) before the lipids were extracted with EtOH/H_2_O to yield a suspension of lipids in EtOH/H_2_O. Centrifugation of the resulting suspension gave a clarified lipid extract which then was concentrated under reduced pressure, desalinated by crystallization in EtOH (abs.), and filtered to give a desalinated lipid concentrate. The final HRO was prepared by standardization and viscosity modification through addition of a reconstituted triglyceride oil and evaporation under reduced pressure.

To generate PEHeRo, lipids were extracted from IHR with EtOH/H_2_O before the resulting extract was clarified by centrifugation and dried under reduced pressure. The dried crude lipid extract was then desalinated by crystallization in EtOH (abs.) and filtration. PEHeRo (>90% PLs w/w dry matter, 85% DM) was purified from the desalinated extract using automatic normal-phase flash column chromatography (EtOH/H_2_O 88:12, 200 nm).

Phospholipid analyses were performed using quantitative ^31^P NMR (Spectral Service AG) and fatty acid analyses were performed using GC-FID according to Ph. Eur. 2.4.29 with modifications to accommodate for PL-rich oils (Mylnefield Lipid Analysis).

### HRO and PEHeRo emulsions

HRO and PEHeRo were prepared as 5% emulsions in dH_2_O by stirring at 800 rpm under inert atmosphere (N_2_ flushing) at 37°C for 1 hour. Emulsions were then pasteurized under inert atmosphere by sonication at 70°C for 2 min in an ultrasonic water bath (Branson 2200, 40 kHz) or by an ultrasonic probe. Emulsions were stored at 4°C for maximum 4 days.

### Macrophage cell culture

Buffy coats were obtained from the blood bank Ålesund (Helse Møre og Romsdal, Norway) with approval by the Regional Ethics Committee (REK, ref.: 230804) and in compliance with the Helsinki Declaration.

Peripheral blood mononuclear cells (PBMC) were isolated from buffy coats by Lymphoprep^TM^ (AXIS-SHIELD PoC AS, 1114547) density gradient and monocytes were selected by plastic adherence and differentiated into monocyte-derived macrophages (MDM) by culture in AIM V medium (Gibco, 12055083) with 5 % CTS™ Immune Cell Serum Replacement (ICSR) and 10 ng/ml M-CSF (Miltenyi Biotec, 130-093-963) for 5 d, followed by incubation without M-CSF.

MDMs from each donor were treated in 1.5 mL per well in a 6-well plate (3 wells per condition) with indicated final percentages (w/w) of HRO or PEHeRo, by dilution of the prepared emulsions in phenol- and serum-free CTS™ AIM-V™ Medium (Gibco, A3830801) supplemented with Pen-Strep. MDMs were either pretreated with HRO for 16 h, followed by change of medium with addition of 1 µg/mL lipopolysaccharide (LPS, Sigma-Aldrich, L2630) and 50 ng/mL IFN-γ (Miltenyi Biotec, 130-096-481) for further 24 h or co-treated with either HRO or PEHeRo and 1 ng/mL LPS for 24 h. Supernatants were collected, pooled per condition, centrifuged and frozen at −80°C until shipping on dry ice for analysis.

For expression analysis of the enzymes required for SPM formation, MDMs were lysed and RNA isolated by the Quick-RNA Miniprep Kit (Zymo Research, R1054). RNA concentration was measured by 260/280 absorbance on a Synergy HTX S1LFA plate reader (BioTek). cDNA was synthesized by the iScript cDNA kit (BioRad, 1708890) and run with the primers for human COX2, ALOX5, ALOX12, ALOX15 and ALOX15B (Supplementary Table S7) and the SsoAdvanced™ Universal SYBR® Green Supermix (BioRad, 1725270) or run with the PrimePCR™ Probe Assays (BioRad) for human HPRT1 (qHsaCIP0030549) and B2M (qHsaCIP0029872) and the SsoAdvanced™ Universal Probes Supermix (BioRad, 1725280) on a CFX96 Real-Time PCR System (BioRad). The mean of HPRT and B2M Cts was used for normalization. Relative mRNA levels were transformed into linear form by the 2 (-ΔΔCt) method. Fold changes of RNA expression relative to unstimulated controls were used for statistical analysis.

### Keratinocyte and fibroblast co-culture

HaCaT keratinocytes (CLS Cell line service GmbH, cat no 300493) were co-cultured with normal human primary dermal fibroblasts (ATCC PCS-201-012) in Dulbecco’s modified Eagle’s medium (DMEM) supplemented with 10% fetal bovine serum (FBS), 100 U/mL penicillin, 100 µg/mL streptomycin and 250 µg/mL fungizone (all purchased from Thermo Fisher Scientific) in T25 tissue culture flasks (Nunc™ EasYFlask™, VWR) and stimulated with and without 100 ng/mL IL-17A (Sigma-Aldrich, cat.no SRP0675) and 5 µg/mL HRO for 96 h, before media were collected and stored at −80 °C for further fatty acid and lipid mediator analysis. HaCaT cells were selected as they represent a commonly used suitable and reliable model for following inflammation and resolution in keratinocytes (63). The co-culture was established by first seeding fibroblasts at 1×10^3^ cells per cm^2^ and pre-cultured in the media described above for 2 days, before adding 4.3 x10^3^ keratinocytes per cm^2^ and co-culturing additional 5 days before initiating stimulation with HRO with and without IL-17A for 96 h as described above. Lower HRO concentrations were used in the keratinocyte/fibroblast experiments than in the MDM experiments due to different viability in the cell models, with the skin cell co-culture being more sensitive towards higher concentrations of HRO than the MDMs.

Total RNA was isolated from treated cells using RNeasy Mini Kit (Qiagen, Hilden, Germany) according to the producers’ protocol. All samples were incubated with RNase Free DNase Set (Qiagen, Hilden, Germany) to remove genomic DNA. RNA concentration and quality were determined using a NanoDrop One Spectrophotometer (Thermo Scientific, Bremen, Germany). LunaScript RT SuperMix Kit (NEW ENGLAND Biolabs) was used to synthesize cDNA following the manufacturer’s protocol. RNA input was 1000 ng per sample. Prior to the qPCR analysis, the specificity of all primers was confirmed by PCR amplification and sequencing (see Supplementary Table S8 for primer information). Briefly, a PCR was run by mixing 2 uL cDNA, 1 μl forward and reverse primer (final concentration of 0.5 μM), 12.5 uL LongAmp Taq DNA polymerase and 8.5 H_2_O. The PCR was run under the following conditions: 95 °C/2min, 35 cycles of 94 °C/20 sec, 60 °C/50 sec, 65 °C/50 sec, followed by 65 °C/10min. The PCR product was then sent to Eurofins Genomics for Sanger sequencing. The qPCR reaction mixture consisted of 4 μl diluted (1:10) cDNA, 1 μl forward and reverse primer (final concentration of 0.5 μM), and 5 μl PowerUp SYBR Green Master Mix (Applied Biosystems, Foster City, California, United States). A standard curve was included for each primer pair to evaluate the primer efficiency. All samples were analyzed in parallels, and non-template and non-enzyme controls were included. The qPCR reaction was run on a QuantStudio 5 instrument (Thermo Fisher Scientific, MA, USA) under the following conditions: Initial denaturation 95 °C for 20 seconds, amplification with 40 cycles at 95 °C for 1 second and 62 °C (60 °C for the reference genes) for 20 seconds, melting at 95 °C for 1 second and 60 °C for 20 seconds, dissociation 95 °C for 1 second. EF1A, RPOL2 and GAPDH were evaluated as reference genes using RefFinder (64) The relative gene expression level was calculated according to the ΔΔCt method with efficiency correction (65) using the geometric averaging of the EF1A, RPOL2 and GAPDH as references.

### Fatty acid composition of cells

Total lipids were extracted from the cells according to the protocol by Folch *et al*. (66). The CHCl_3_ phase was dried under nitrogen gas, and thereafter the samples were methylated overnight with benzene, 2% H_2_SO_4_ in MeOH, and dimethoxypropanate at 50 °C. The methyl esters of the fatty acids were separated using GC (Hewlett Packard 6890) with flame-ionizing detector (FID) using a BPX70 column (60 m, 0.25 mm i.d., 0.25 µm film) with a temperature program starting at 50 °C, and increasing at a rate of 4 °C /min until 170 °C, then raised to 0.5 °C/min until 200 °C, and finally to 10 °C/min until 240 °C. The peaks were integrated using GC ChemStation software (Agilent Technologies, Rev. B.01.01 SR1, 2001-2005). The fatty acids were identified by an external standard (GLC 463, Nu-Chek Prep), and the concentration of the individual fatty acids was calculated based on an internal standard (C23:0, Larodan). For analysis of fatty acid composition in lipid classes, the CHCl_3_ lipid extracts were first evaporated under nitrogen gas and re-dissolved in hexane. The lipid classes (phospholipids, triacylglycerides and free fatty acids) were separated by thin layer chromatography using a mixture of petroleum ether, Et_2_O and AcOH (113:20:2, by volume) as the mobile phase. The lipids were visualized by spraying the TLC plates with 0.2% (w/v) 2′,7′-dichlorofluorescein in MeOH, and then identified by comparing with known standards (Sigma Chemical Co.) under UV light. The spots corresponding to the different fractions were scraped off into glass tubes, trans-methylated following the aforementioned procedure and the fatty acid composition determined by GC-FID.

### Lipid mediator analyses

Lipid mediator analysis was performed using methodologies and approaches recently published by Dooley *et al.* (67).

Snap frozen media was placed in 2 volumes of MeOH containing deuterated internal standards (IS) (d_4_-LTB_4_, d_5_-MaR1, d_5_-MaR2, d_4_-PGE_2_, d_5_-LXA_4_, d_5_-RvD_3_, d_5_-RvD2, d_4_-RvE1, d_5_-17R-RvD1, d_5_-LTE_4_, d_5_-LTD_4_ and d_5_-LTC_4_), representing the chromatographic regions of interest, were added to facilitate lipid mediator identification and quantification. This was placed at −20 °C for at least 45 minutes then centrifuged and supernatants were subjected to solid-phase extraction (SPE) using the ExtraHera system (Biotage) and Isolute C18 500 mg columns (Biotage).

Oils samples were placed in 2 volumes of MeOH containing 2.5 ng of deuterated IS (d_4_-LTB_4_, d_5_-MaR1, d_5_-MaR2, d_4_-PGE_2_, d_5_-LXA_4_, d_5_-RvD_3_, d_5_-RvD2, d_4_-RvE1, d_5_-17R-RvD1, d_5_-LTE_4_, d_5_-LTD_4_ and d_5_-LTC_4_) and incubated in 2 volumes of KOH (1N) for 45 minutes at 45 °C. At the end of the incubation the samples were immediately acidified and subjected to SPE using the ExtraHera system (Biotage) and Isolute C18 500 mg columns (Biotage). Oils analysed for unesterified lipid mediators were not saponified prior to being subjected to SPE.

Methyl formate and MeOH fractions from C18 SPE were collected, brought to dryness, and suspended in phase (MeOH/H_2_O, 1:1. v/v) for injection on a Shimadzu LC-20AD HPLC and a Shimadzu SIL-20AC autoinjector, paired with a QTrap 6500+ (Sciex). Analysis of mediators isolated in the methyl formate fraction was conducted as follows: an Agilent Poroshell 120 EC-C18 column (100[mm[×[4.6[mm[×[2.7[µm) was kept at 50 °C and mediators eluted using a mobile phase consisting of MeOH/H_2_O/AcOH of 20:80:0.01 (v/v/v) that was ramped to 50:50:0.01 (v/v/v) over 0.5 min and then to 80:20:0.01 (v/v/v) from 2 min to 11[min, maintained till 14.5 min and then rapidly ramped to 98:2:0.01 (v/v/v) for the next 0.1 min. This was subsequently maintained at 98:2:0.01 (v/v/v) for 5.4 min, and the flow rate was maintained at 0.5[mL/min. In the analysis of mediators isolated in the MeOH fraction, the initial mobile phase was MeOH/H_2_O/AcOH of 20:80:0.5 (v/v/v) which was ramped to 55:45:0.5 (v/v/v) over 0.2 min and then to 70:30:0.5 (v/v/v) over 5 min and then ramped to 80:20:0.5 (v/v/v) for the next 2 min. The mobile phase was maintained for 3 min and ramped to 98:2:0.5 (v/v/v) for 2 min. QTrap 6500+ was operated using a multiple reaction monitoring (MRM) method. Each lipid mediator was identified using the following criteria: (1) matching retention time to synthetic or authentic standards (±0.05 min), (2) a signal-to-noise ratio ≥ 5 for a primary transition, and (3) a signal-to-noise ratio of ≥3 for a secondary transition. Data were analyzed using validated protocols (67) and Sciex OS v3.0. Chromatograms were reviewed using the AutoPeak algorithm, using ‘low’ smoothing setting and signal to noise ratios were calculated using the relative noise algorithm (67). External calibration curves were used to quantify identified mediators. Where available calibration curves were obtained for each mediator using synthetic compound mixtures that gave linear calibration curves with R^2^ values of 0.98–0.99. These calibration curves were then used to calculate the abundance of each mediator. Where synthetic standards were not available for the construction of calibration curves, calibration curves for mediators with similar physical properties (e.g. carbon chain length, number of double bonds, number of hydroxyl groups and similar elution time) were used.

### Statistical analysis

Statistical significances were determined using; the Mann-Whitney U test (one-tailed/two-tailed as appropriate) for comparison of two groups (except for the fatty acid composition experiments in Figure 2) and one-factor ANOVA for comparison of >2 groups. Post[hoc comparisons included Tukey’s HSD or Dunnett[type comparisons with Holm familywise error rate adjustment. Statistical significance criteria for difference between groups were p < 0.05. All results are presented as mean values ± the standard error of the mean (SEM). Statistical analysis of fatty acid composition in cells were performed using Unistat v10 (Unistat Ltd., London, UK), PLS-DA analyses were performed using MetaboAnalyst v6.0 (68), analyses of fold changes of RNA expression were performed using GraphPad Prism v10.6.1, all other statistical analyses were performed using Python (Python version ≥3.10) with the libraries statsmodels, scipy.stats, and numpy (scripts can be found here: https://github.com/tringheimbakka/IPN327953).

## Results

### Oil composition and lipid mediator content in HRO and PEHeRo

To understand the effects of the phospholipid ester rich oils in cellular systems, it was important to understand the composition of the oils both with regards to fatty acid profile and lipid class distribution, but also with regards to lipid mediators potentially present in the oil. We therefore began with characterizing the composition of the oils.

Herring roe oil (HRO) is a lipid extract from herring roe that has been standardized with regard to content of *phospholipid esters from herring roe* (PEHeRo) using a marine triacylglyceride (TAG) oil. The phospholipid esters in PEHeRo are predominantly phosphatidylcholine (>80%) and enriched with LC-PUFAs. The fatty acid profiles in both HRO and PEHeRo display a high content of DHA with a DHA/EPA ratio of approximately 3:1, where DHA and EPA make up 50-60% of the total fatty acid profile in area %. The HRO and PEHeRo investigated in this work had a phospholipid content of 34% and 91% (per anhydrous substance) and 425 mgTAG/g and 318 mgTAG/g DHA+EPA, respectively. Lastly, HRO and PEHeRo were found to contain 6% area and 1% area of omega-3 docosapentaenoic acid (n-3 DPA), respectively.

After establishing the macroscopic lipid composition, the content of lipid mediators in HRO and PEHeRo was determined, as it is possible that the lipid mediator levels in the oils could influence their behavior and resulting SPM biosynthesis in the cell systems. The oils were screened for lipid mediators, including SPMs, originating from DHA, n-3 DPA, EPA, and arachidonic acid (AA) and the full results are presented in Supplementary Table S1. The qualitative lipid mediator profiles of the two oils were, unsurprisingly, found to be similar, with the predominant SPMs originating mostly from the DHA metabolome in the form of RvD5, RvD6, PDX, 17R-PD1, MaR1, and MaR2 (Fig. 1A). Furthermore, there were also notable contributions from SPMs originating from the n-3 DPA (RvT4 and RvD5 _n-3 DPA_) and EPA (RvE4) metabolomes, whereas lipid mediators from the AA metabolome, as expected, only made up 1-2% of the total SPM content in the oils.

**Fig. 1.**
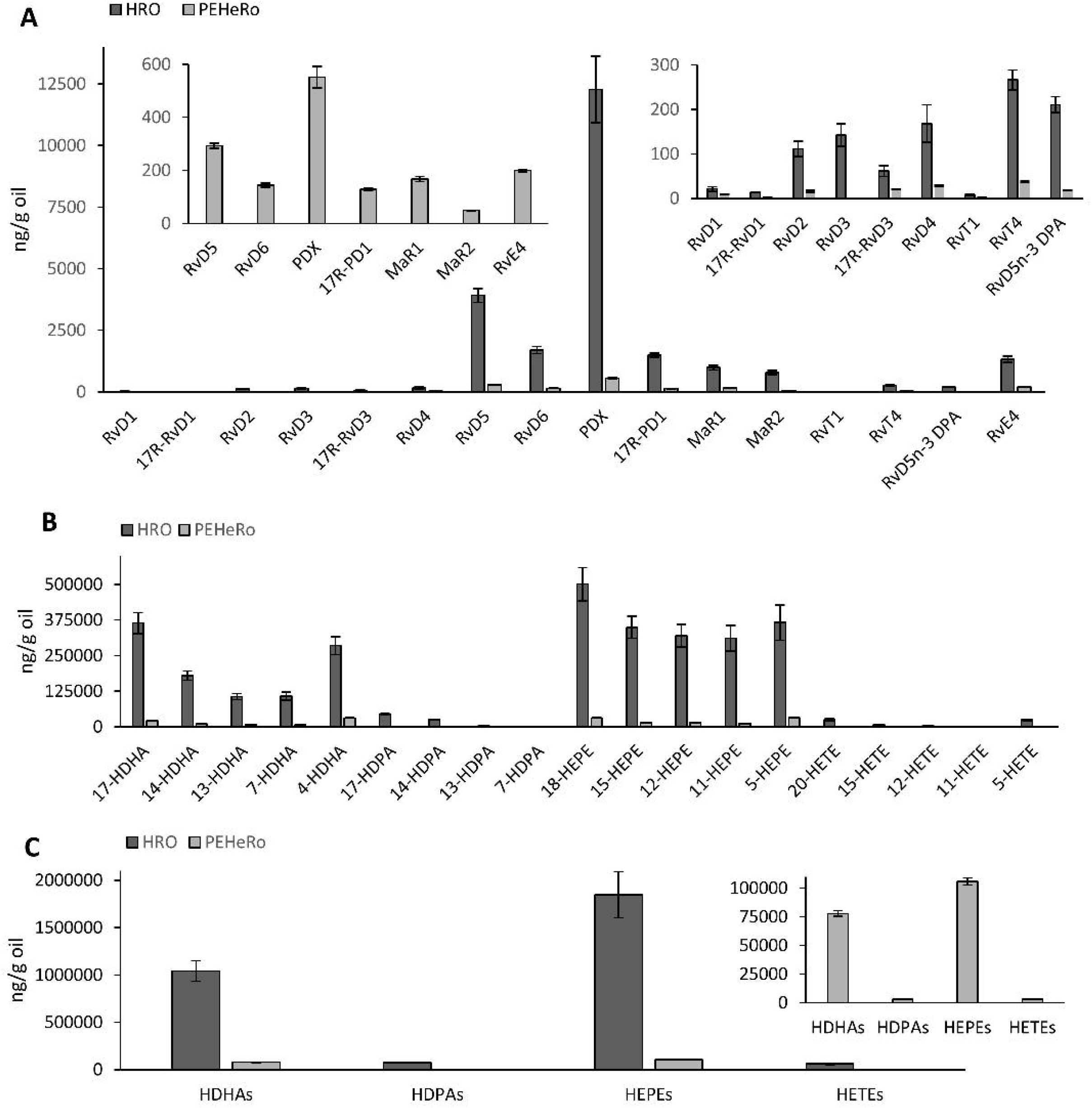
A) SPMs in HRO and PEHeRo (ng/g oil, mean of 3 replicates ±SEM). B) Monohydroxylated fatty acids in HRO and PEHeRo (ng/g oil, mean of 3 replicates ±SEM) C) Sum of monohydroxylated fatty acids in HRO and PEHeRo from DHA, n-3 DPA, EPA, and AA (ng/g oil, mean of 3 replicates ±SEM). Lipid mediators and SPMs were identified and quantified using LC-MS/MS.

**Fig. 2.**
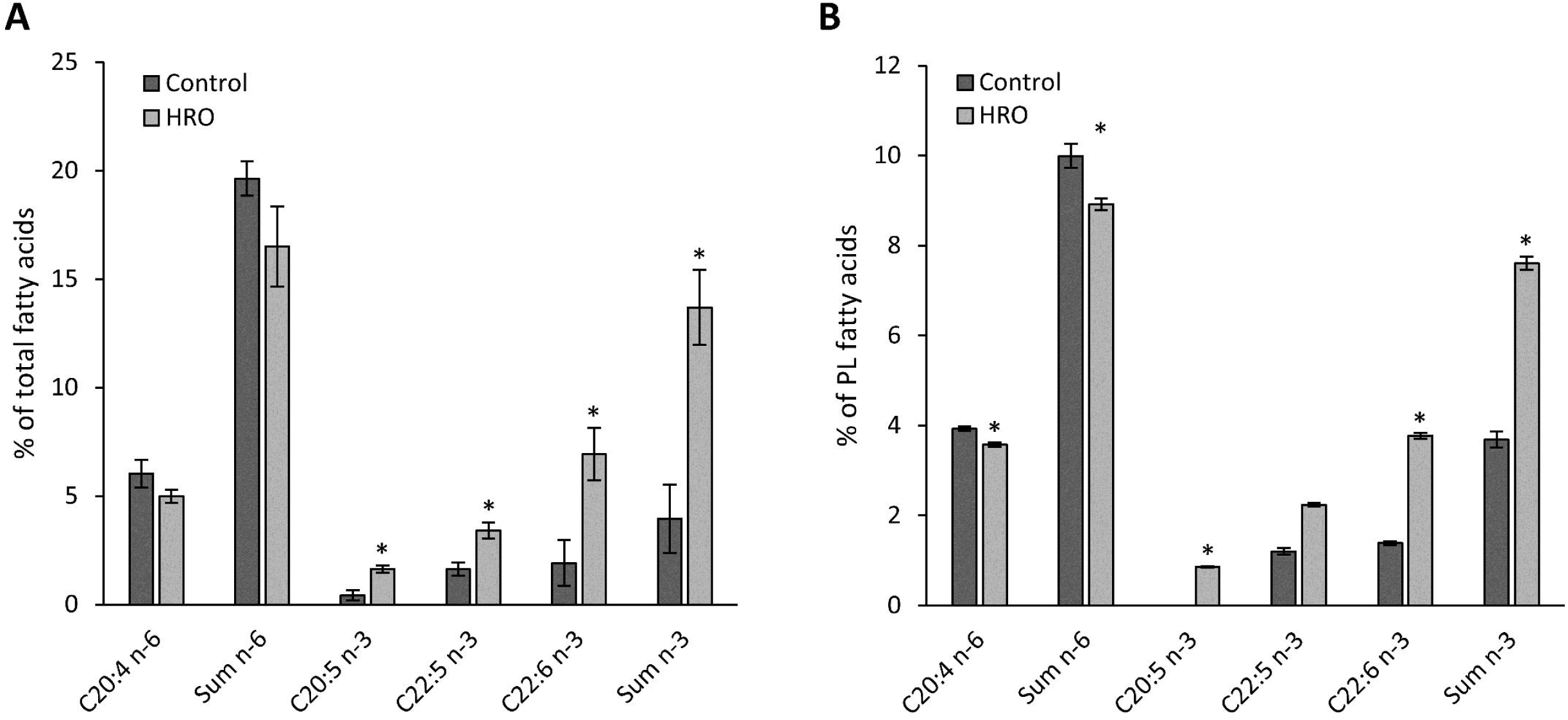
A) Relative concentrations from GC-FID of fatty acids as % of total fatty acids in MDMs (mean of 3 replicates ±SEM) when treated with HRO versus control. B) Relative concentrations from GC-FID of fatty acids as % of total fatty acids in the phospholipid fraction of the keratinocyte/fibroblast co-culture (mean of 9 replicates ±SEM) when treated with HRO versus control. *) p < 0.05 (One-factor ANOVA with Tukey’s HSD).

The key difference between HRO and PEHeRo regarding lipid mediator content, was the >10-fold higher concentration of lipid mediators in the less refined HRO compared to the concentrated phospholipid oil PEHeRo. The levels of SPMs from the DHA, n-3 DPA, and EPA metabolomes in PEHeRo were 6% - 15% of the concentration in HRO, and monohydroxylated fatty acids at 4% - 7% of the levels observed in HRO.

The levels of monohydroxylated fatty acids (HDHAs, HEPEs, HDPAs, and HETEs) were found to be surprisingly high in HEPEs given the overall fatty acid composition, with 5-HEPE and 18-HEPE being the two major contributors (Fig. 1B and Fig. 1C). The HEPE/HDHA ratio was found to be 1.8 and 1.4 for HRO and PEHeRo respectively (Fig. 1C), where a significant portion (>50%) of total HEPEs and HDHAs were found to be present as metabolically available free fatty acids rather than bound in lipids. The sum of HEPEs and HDHAs made up >95% of the total amount of monohydroxylated fatty acids in the two oils. Lastly, higher levels of unesterified EPA were observed in both HRO and PEHeRo compared to DHA free fatty acid levels.

### Supplemented HRO is taken up by macrophages and keratinocytes/fibroblasts

The uptake of fatty acids from HRO was analyzed by GC-FID, showing a significant increase in DHA, n-3 DPA, and EPA in total lipids in the MDMs (Fig. 2A and Supplementary Table S2). In the direct keratinocyte/fibroblast co-culture, it was further shown that the changes introduced by HRO could be observed in the phospholipid fractions of the cells with similarly significantly increased EPA and DHA and decreased AA (Fig. 2B and Supplementary Table S3).

### HRO and PEHeRo promote SPM biosynthesis in monocyte-derived macrophages

The macrophage experiments conducted by Mildenberger *et al.* demonstrated reduction in cytokine secretion, and we therefore wanted to study SPM biosynthesis in the macrophages treated with HRO in order to better understand how this oil and its phospholipids (PEHeRo) elicit their anti-inflammatory and possibly pro-resolving activities (60).

We therefore quantified the SPMs present in the supernatants from likewise stimulated MDMs, which showed a significant upregulation of SPM concentrations when macrophages were pre-treated with HRO prior to stimulation with LPS+IFN-γ (Fig. 3A-D and Supplementary Table S4). The lipid mediator biosynthesis was found to be significantly upregulated for SPMs from the DHA, EPA, and AA metabolomes, including a non-significant trend for upregulation of n-3 DPA SPMs. The EPA-derived SPM RvE3 was found to have the highest concentration in the treated samples. The overall SPM profiles of the treated macrophages showed activation of both ALOX5 and ALOX15 pathways.

**Fig. 3.**
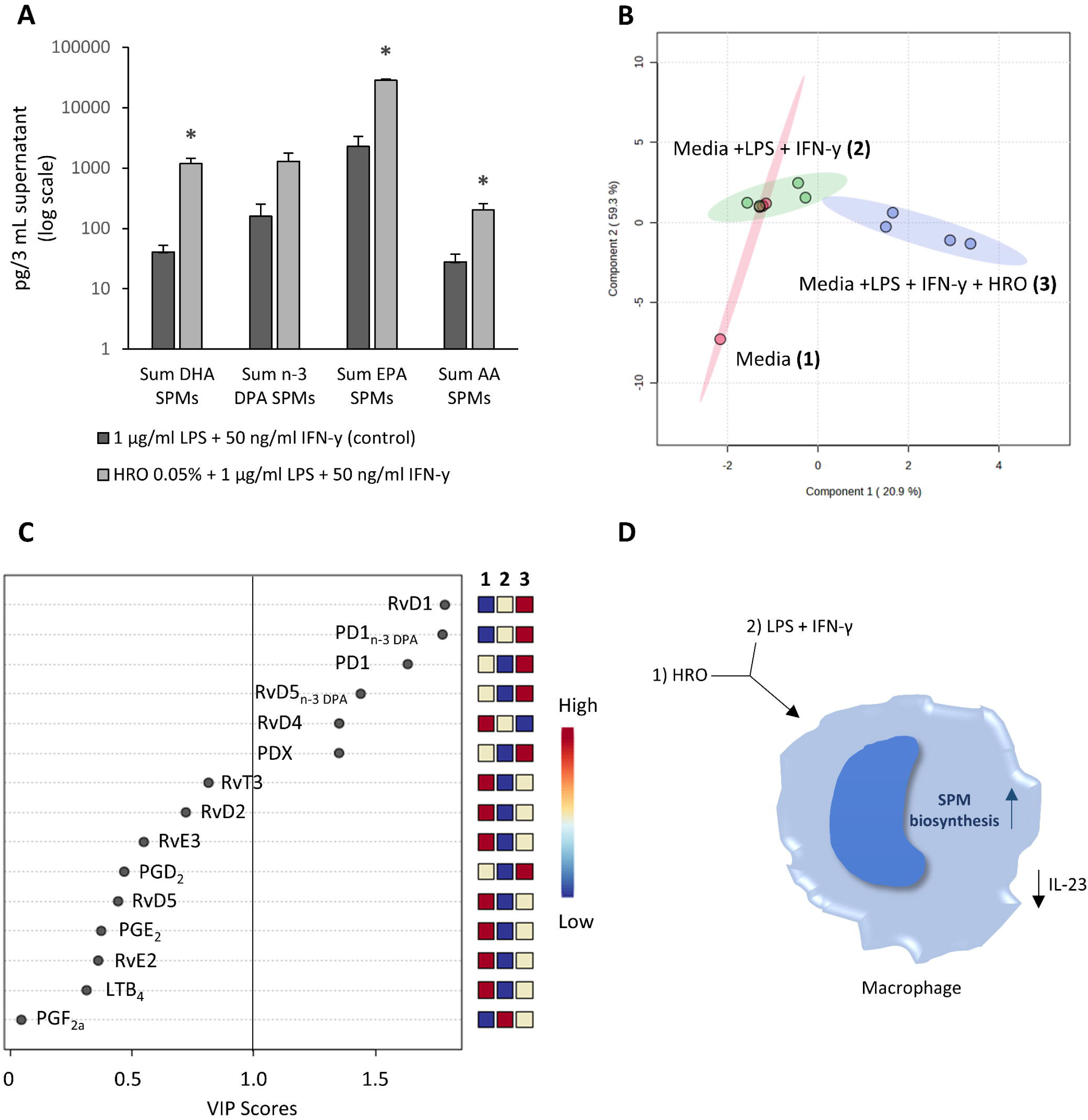
A) SPM levels (mean of 4 donors/experiments ± SEM) in supernatants from macrophages incubated with HRO prior to stimulation with LPS+IFN-γ versus only stimulated with LPS+IFN-γ (control). B) Partial least squares discriminant analysis (PLS-DA) of supernatants from macrophages incubated with HRO prior to stimulation with LPS+IFN-γ versus only stimulated with LPS+IFN-γ (control). Results are compared to lipid mediator concentrations identified in fresh media. C) Variable importance in projection (VIP) scores of SPMs that contribute to the difference between the tested conditions. D) Figure showing the treatment conditions and the highlighted measured outcomes. SPMs were identified and quantified using LC-MS/MS. *) p < 0.05 (Mann-Whitney U test, two-tailed).

Partial least squares discriminant analysis (PLS-DA) of the lipid mediators in the cell supernatants from the treated and untreated macrophages (including a fresh media control sample) as shown in Fig. 3B also displayed a difference between the groups, where the shifts in SPM concentrations are visualized by the separation of the clusters in green (2, LPS+IFN-γ) and blue (3, LPS+IFN-γ+HRO) in the PLS-DA scores plot. The chart shown in Fig. 3C shows the variable importance in projection (VIP) scores of individual SPMs that contribute to the separation of the clusters in the PLS-DA with a VIP score >1 identifying those mediators that contribute to the observed separation to the greatest extent. VIP scores >1 were observed for four SPMs from the DHA metabolome (RvD1, RvD4, PD1, and PDX) and two SPMs from the n-3 DPA metabolome (RvD5_n-3_ DPA and PD1n-3 DPA).

Considering the promotion of SPM biosynthesis observed in the macrophages when pretreated with HRO, we wanted to further expand upon this and see if we were able to optimize the experiment setup with regards to detecting and measuring SPMs in the supernatants after treatment/stimulation. Also, considering that the initial experiments were performed on the entire HRO, we also wanted to evaluate if there were any differences between HRO and a purified sample of PEHeRo in terms of ability to promote lipid mediator biosynthesis in the primary immune cells. So, we tested two concentrations of HRO (0.05% and 0.25%) and one concentration of PEHeRo (0.25%) in an experiment setup where the macrophages were co-stimulated with the oils and LPS (1.0 ng/mL) for 24 hours, similarly to the conditions employed by Sobrino *et al.* (37).

Similarly to the initial experiment, we also observed a significant upregulation of SPM biosynthesis in the macrophages treated with HRO and PEHeRo (Fig. 4A-D and Supplementary Table S5). The results also showed, as expected, generally higher concentrations of SPMs in the cell supernatants, thus offering the potential for greater understanding of the pathways activated upon treatment with HRO and PEHeRo.

**Fig. 4.**
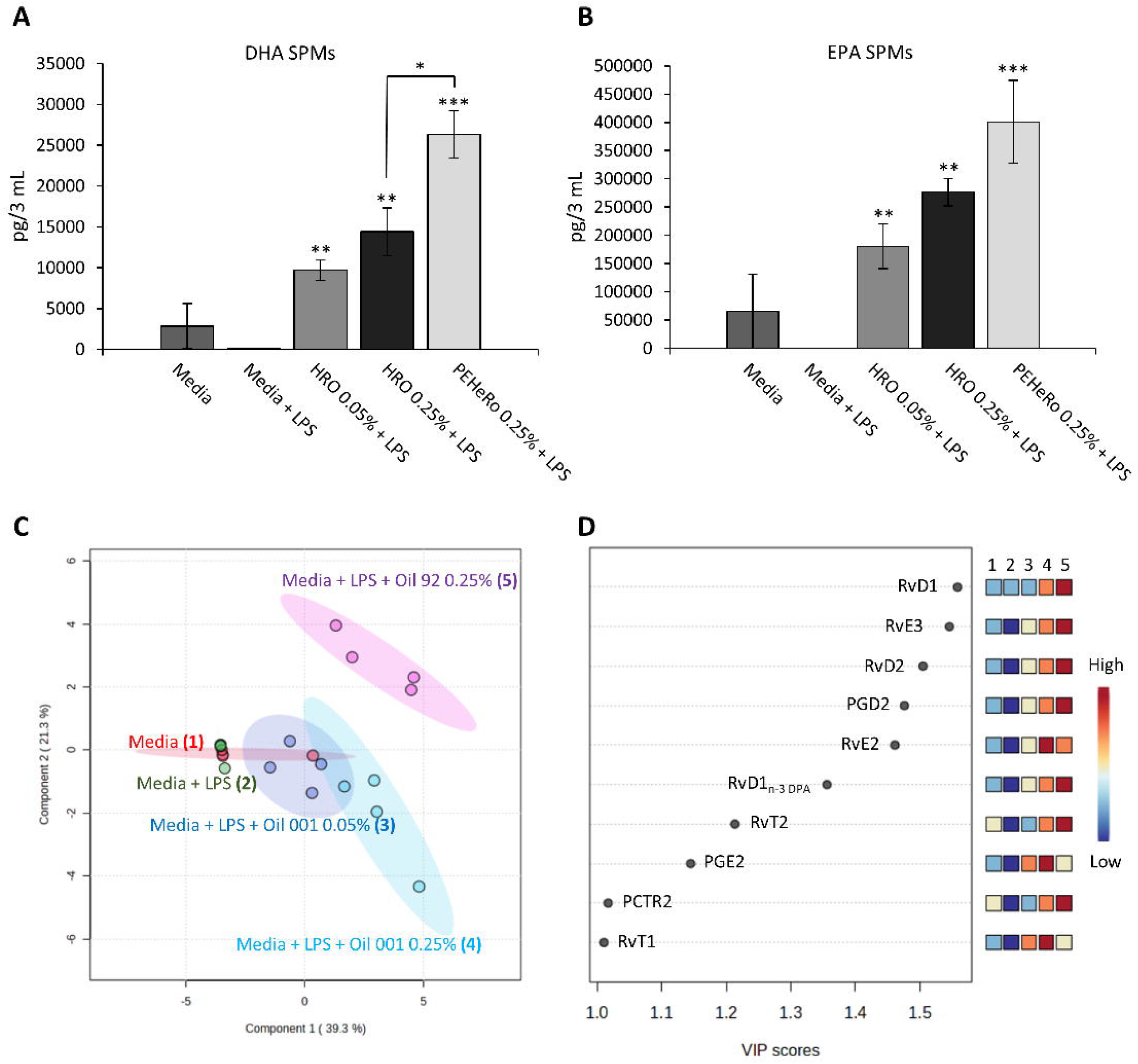
SPM levels (mean of 4 donors/experiments ± SEM) from DHA (A) and EPA (B) metabolomes in macrophage cell supernatants (pg/3 mL) when incubated with HRO+LPS, PEHeRo+LPS, LPS (control), or untreated. C) Partial least squares discriminant analysis (PLS-DA) of supernatants from macrophages incubated with HRO+LPS, PEHeRo+LPS, LPS (control), or untreated. D) Variable importance in projection (VIP) scores of SPMs that contribute to the difference between the tested conditions. SPMs were identified and quantified using LC-MS/MS. *) p < 0.05, **) p < 0.01, and ***) p < 0.001 (One-factor ANOVA with one-way Dunnett-type comparison for comparison to control and Tukey’s HSD for comparison between treatment groups).

All three treatment conditions (0.05% HRO, 0.25% HRO, and 0.25% PEHeRo) gave significant upregulation of the sum of SPMs from the DHA and EPA metabolomes (Fig. 4A-B) when compared to media stimulated with LPS (control). We also observed a non-significant trend for upregulation of SPMs from the DPA metabolome and, in contrast to the first experiment, no upregulation of AA SPMs. Similarly to the first experiments, the most strongly expressed SPM was the EPA-derived SPM RvE3. Furthermore, when comparing the two treatment concentrations of HRO (0.05% and 0.25%), we observed a non-significant dose-dependent trend where the higher treatment concentration gave higher mean values compared to the lower concentration. The macrophage SPM biosynthetic response to the treatment did not reflect the 5-fold increase in treatment concentration and may indicate saturation of the system with regards to accessible fatty acids for SPM biosynthesis.

Looking at individual SPMs through the PLS-DA analysis from the experiments, it was observed that PEHeRo elicited a different response from the treated macrophages compared to the two concentrations of HRO. This is well visualized through the clear separation of the cluster for PEHeRo and the two HRO clusters in Fig. 4C. The partial overlap of the two different doses of HRO was in alignment with the observation made for the abundance of SPMs from the different metabolomes, where a non-significant trend for dose-dependency was observed (Fig. 4A-B). The VIP scores shown in Fig. 4D indicate which individual SPMs that are drivers for the separation of the clusters similarly to Fig. 3C. The VIP scoring shows also here that RvD1 was the biggest driver for separation of the clusters, followed by RvE3 which was found to be the most abundant SPM present in both experiments across all concentrations. Furthermore, we observed that individual SPMs from all four metabolomes (DHA, n-3 DPA, EPA, and AA) contributed to the separation of the clusters. Despite not observing significant upregulation of total n-3 DPA SPMs between the groups, the biosynthesis of the following individual n-3 DPA SPMs with a VIP-score >1 were found to be significantly upregulated: RvD1 _n-3 DPA_ for PEHeRo at 0.25%, RvT2 for PEHeRo at 0.25%, and RvT1 for both concentrations of HRO. These observations corroborate the significance of n-3 DPA SPMs for the separation of the PLS-DA clusters.

Interestingly, the VIP scores in Fig. 4D also showed the cysteinyl DHA-derived SPM PCTR2 (PCTR = protectin conjugates in tissue regeneration) from the DHA metabolome to be a significant contributor to the separation of the different clusters, thus indicating activation of reparative pathways and not only protective SPM pathways.

To support the hypothesis of the observed SPMs and lipid mediators arose from upregulated biosynthesis in the macrophages, we assessed the gene expression of key biosynthetic enzymes following treatment with HRO, LPS, and IFNγ using RT-qPCR. The enzymes COX2, ALOX5, ALOX12, ALOX15, and ALOX15B were selected as they represent a large fraction of the central enzymes involved lipid mediator biosynthesis (10,69). RT-qPCR showed that all enzymes were expressed except ALOX15, which was not detected (Fig. S1). These findings support the hypothesis that the lipid mediators observed in the supernatants were indeed produced by the macrophages. In addition to the absence of ALOX15 gene expression and detection of ALOX15B, an interesting observation was that HRO appeared to attenuate COX[2 gene expression in macrophages.

### HRO promotes SPM biosynthesis in IL-17A stimulated fibroblast/keratinocyte co-culture

To study the potential effects of HRO on the IL-23/IL-17 cytokine axis in psoriasis, a co-culture of keratinocytes and fibroblasts that mimics psoriasis upon stimulation with IL-17A was developed. Mildenberger *et al.* (60) showed that HRO dampened psoriatic marker S100A7 (psoriasine) and the IL-17 response gene NFΚBIZ in this co-culture. Considering the dampening effects observed in these co-culture experiments, we wanted to see if the SPM biosynthesis was upregulated in the co-culture.

Quantification of SPMs and monohydroxylated fatty acids by LC-MS/MS in the co-culture cell media showed significant upregulation of SPMs from the DHA and EPA metabolomes, as well as significant upregulation of monohydroxylated DHA, EPA, and n-3 DPA (Fig. 5A-D and Supplementary Table S6). It was interesting to observe a significant promotion of SPM biosynthesis taking place despite only treating the cells with 1/100 (0.0005%) of the HRO amount compared to the 0.05% HRO concentration used in the macrophages, as well as reporting on only 1/3 of the media volume (3 mL vs 1 mL). The lipid mediator levels measured in the media were affected by the decreased concentration of HRO, and the overall quantities were lower compared to what was observed in the macrophage experiments.

**Fig. 5.**
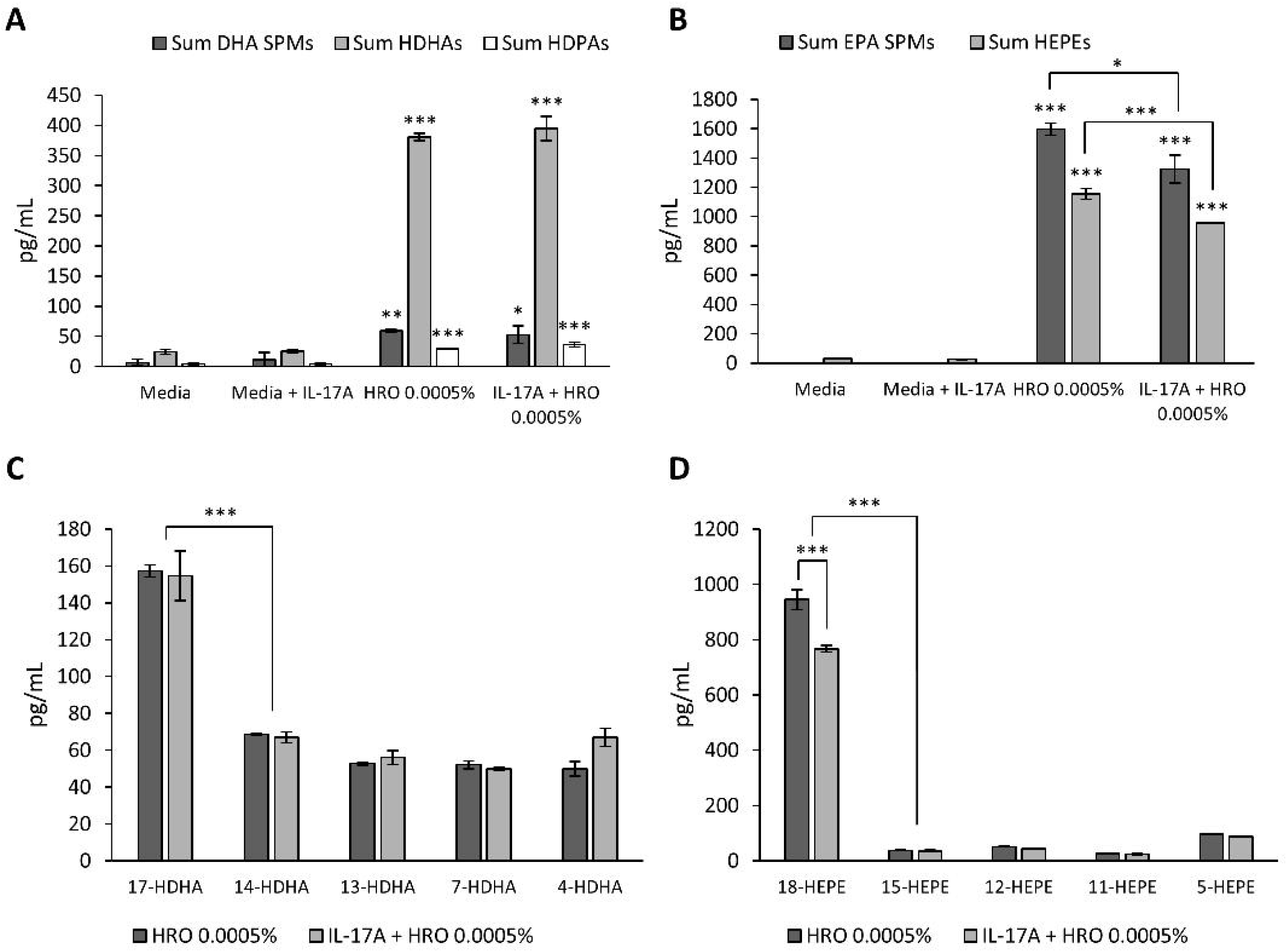
SPM and SPM precursor levels (mean of 3 biological replicates ± SEM) in fibroblast/keratinocyte co-culture cell supernatants (pg/mL) incubated with IL-17A (control), HRO, HRO+IL-17A, or untreated. SPMs were identified and quantified using LC-MS/MS. A) SPMs and monohydroxylated fatty acids from DHA and monohydroxylated fatty acids from DPA. B) SPMs and monohydroxylated fatty acids from EPA. C) Monohydroxylated fatty acids from DHA. D) Monohydroxylated fatty acids from EPA. *) p < 0.05, **) p < 0.01, and ***) p < 0.001 (One-factor ANOVA with one-way Dunnett-type comparison for comparison to control and Tukey’s HSD for comparison between treatment groups and between different monohydroxylated fatty acids).

The DHA SPMs that contributed to the significant upregulation of sum of DHA SPMs (Fig. 5A) in both the groups treated with HRO included RvD1, RvD2, RvD3, RvD5, RvD6, and PD1, of which RvD5 and RvD6 were significant in HRO+IL-17A compared to the stimulated control. The contributors to the overall activation of the DHA SPM metabolome coincide with observations made in the macrophage experiments where RvD1 and PD1 had VIP scores >1 when HRO was used to pretreat macrophages prior to stimulation with LPS+IFN-γ and RvD1 and RvD2 had VIP scores >1 when macrophages were co-stimulated with HRO/PEHeRo + LPS.

The significant upregulation of monohydroxylated fatty acids (HDHAs and HDPAs) could also support that the HRO indeed promoted activation of the metabolic pathways for SPM biosynthesis from the DHA and DPA metabolomes. The HDHA profile measured in the supernatants supported the observations made for the SPMs (Fig. 5C), as the main monohydroxylated fatty acid was 17-HDHA, where the 17S-HDHA isomer is a precursor for the D-series resolvins formed through ALOX15. 17-HDHA was significantly favored over 14-HDHA with the 14S isomer being a pathway marker for the biosynthesis of MaR1 and MaR2 through ALOX12. The apparent preference for 17-HDHA thus corresponded with the observed levels of MaR1 and MaR2 where neither were measured above the LLOQ. The same was the case for the experiments with the macrophages where no consistent or significant observations of MaR1 and MaR2 were made. It should be noted that this apparent pathway domination was more pronounced for the HDPAs, as only 17-HDPA was measured in the samples.

Furthermore, SPMs from the EPA metabolome were also significantly increased in both HRO treatment groups compared to the stimulated control (Fig. 5B). This was primarily driven by RvE3 and to a lesser extent RvE2 (approximately 1:40 RvE2/RvE3), both of which were also separately significantly upregulated compared to the stimulated control. RvE2 and RvE3 being the main EPA SPMs measured was not surprising when looking at the HEPE profile of both treatment groups (Fig. 5D), where 18-HEPE was found in the, by far, highest concentration in the sample. The 18R isomer of 18-HEPE is a SPM precursor biosynthesized via CYP450 and GPX4 and is the main intermediate before a one-step conversion via ALOX15 into RvE3 or the conversion to RvE2 via ALOX5. The abundance of 15-HEPE was far lower than the abundance of 18-HEPE, which may explain why RvE4 was not found above the LLOQ as 15S-HEPE is the main precursor for RvE4.

When comparing HRO treated cells with and without IL-17A stimulation, it was observed that the IL-17A stimulation itself might have impacted lipid mediator biosynthesis. The greatest impact of IL-17A co-stimulation was observed for the EPA metabolome, with a significant reduction of 18-HEPE, sum HEPEs, and EPA SPMs when comparing the HRO group with the HRO+IL-17A group. Interestingly, only RvE3 was significantly reduced, whereas only a non-significant trend was observed for RvE2. An impact of IL-17A stimulation was also observed for SPMs in the DHA metabolome, albeit to a lesser extent. Here we observed that both RvD2 and PD1 were significantly upregulated in the HRO group, but not in the HRO+IL-17A group.

Gene expression of selected enzymes was also assessed by RT-qPCR in the skin cell co-culture. Treatment with HRO produced minimal changes in gene expression relative to control (Fig. S2). However, transcripts for COX2, ALOX15, ALOX15B, ALOX12, and 15-LOX were all detectable under the treatment conditions.

Thus, showing that the full enzymatic machinery required for lipid mediator synthesis was present in the skin cell co-culture model.

## Discussion

An interesting finding from the macrophage experiments in the present study was the comparison of total SPMs across different metabolomes. Where notably, treatment with PEHeRo significantly increased the biosynthesis of SPMs from the DHA metabolome compared to HRO. In addition to the significant difference between PEHeRo and HRO in the DHA metabolome, a similar observation was made for SPMs from the EPA metabolome albeit as a non-significant trend. The observed increase in SPM biosynthesis in macrophages treated with PEHeRo cannot be explained by availability of metabolically available free fatty acid precursors or availability of mono-hydroxylated fatty acids, including lipid mediator precursors, as PEHeRo was shown to contain far less of these compounds compared to the HRO used in these experiments. Thus, any normalization based on availability of free EPA/DHA, bound EPA/DHA, or monohydroxylated fatty acids (18-HEPE and 17-HDHA) only increased the difference observed when comparing HRO and PEHeRo based on overall concentration. This indicated that the more purified PEHeRo more efficiently promoted upregulation of biosynthesis and that the oils’ ability to upregulate these pathways depends on other factors such as matrix complexity or lipid class composition and not just the precursor availability. For instance, phospholipids have been indicated to give increased *in vivo* bioavailability of PUFAs compared to corresponding triacylglycerides (49,70), and their lysoforms may also themselves act as signaling molecules (71). The observations correspond with existing data that indicates that different oils elicit different ability to promote conversion of PUFAs and SPM precursors into SPMs (37). Pinpointing the impacting factors for these observations needs further studies to be verified.

Consistent dominant upregulation of the EPA-derived SPMs RvE2 and RvE3 was observed in all experiments, with RvE3 being the most abundant of the two. The higher relative concentration of RvE3 compared to other SPMs can possibly be attributed to RvE3 being available in one enzymatic transformation from 18R-HEPE by ALOX15B, which was likely abundant in both HRO and PEHeRo from the measured levels of 18-HEPE (achiral analysis). RvE2 is also biosynthesized from 18R-HEPE but through two steps by ALOX5 and a dehydrogenase. The availability of 18-HEPE in the oils does not fully explain the strong upregulation of RvE3, as more RvE3 was found in cell media when the macrophages were stimulated with PEHeRo which contained >10-fold less 18-HEPE than HRO. The consistent upregulation of RvE2 and RvE3 across both the macrophages and the skin cell co-culture experiments was particularly interesting given that RvE2 has been reported to be a regulator of neutrophil infiltration in addition to enhancing phagocytosis (72), and more importantly, RvE3 has been reported to attenuate inflammation via the IL-23/IL-17A-axis (73) which is highly relevant for psoriatic inflammation.

The most recurring DHA-derived SPM was found to be RvD2, which was observed consistently in the macrophage experiments and for the HRO treated group in the keratinocyte/fibroblast co-culture experiments. RvD2 takes part in regulating inflammation by increasing phagocytosis and reducing neutrophil infiltration (74), promoting macrophage polarization (75), and reducing expression of proinflammatory cytokines (74).

The protectin PDX has been reported to be implicated in regulation of psoriasis, as it has been found to be elevated in psoriasis lesional skin (76), and was measured in all the macrophage experiments. By being upregulated in the immune cells treated with HRO and PEHeRo and involved in regulation of inflammation in psoriasis, PDX could be a promising biomarker for future studies in psoriasis using marine phospholipids.

Showing that we were able to upregulate SPM biosynthesis in the keratinocyte/fibroblast co-culture was intriguing, considering that there, to our knowledge, are few studies detailing the ability of lipid supplementation to regulate SPM production in these cells. The monohydroxylated fatty acid profiles could imply that a key pathway for synthesis of lipid mediators from DHA and DPA goes through 17-HDHA, as there were found significantly higher concentrations of 17-HDHA compared to 14-HDHA. This was even more pronounced for n-3 DPA, where only 17-HDPA was found in the media upon analysis. Also, it was interesting to see activation of the DPA metabolome considering the much lower content of this PUFA in HRO, thus implicating the importance of this pathway in regulation of inflammation. The apparent preference for the 17-hydroxylated DHA and n-3 DPA are aligned with literature showing 17-HDHA biosynthesis through ALOX15 in normal human keratinocytes when treated with DHA (77). Morin *et al.* have shown that supplementation of a psoriatic skin cell model with pure EPA also led to increased epidermal production of 17-HDHA, besides PGE_3_ and 12-HEPE, but not in the presence of polarized T-cells (78). Future experiments in the keratinocyte/fibroblast co-cultures should include chiral analyses of the monohydroxylated fatty acids to further elucidate which enzymes that are involved. Our findings furthermore indicate that the SPM biosynthesis from DHA and DPA is maintained in an inflamed state, as the precursor levels were found to be very similar between the HRO and the HRO + IL-17A groups. However, where ALOX15 seems to be important for conversion of DHA and DPA, this does not seem to be the case for EPA. Biosynthesis via 18-HEPE seems to be the main pathway for EPA derived lipid mediators with 15-HEPE levels being comparatively lower. This could be due to substrate vs enzyme availability in the system, but the finding is also consistent with the SPMs found in both the skin cells and the macrophages, where RvE4, a downstream product of 15S-HEPE, was not measured in any of the experiments.

The observed biosynthesis of the cysteinyl SPM PCTR2 in macrophages may also indicate a shift towards a reparative phenotype as well as a protective one, as the PCTRs have been reported to be biosynthesized by macrophages and to be involved in macrophage phenotype regulation and tissue regeneration (12,79,80). The PCTRs are biosynthesized similarly to PD1 through ALOX15 conversion of 17S-HpDHA into 16S,17S-epoxy-DHA, but with a last step through a glutamyltransferase instead of an epoxide hydrolase to give PCTR1 instead of PD1. PCTR1 is then hydrolyzed to PCTR2 which may be taken through a final hydrolysis to PCTR3. The linear nature of their biosynthesis implies that PCTR1 also was produced by these cells prior to being hydrolyzed to PCTR2, which is interesting given that PCTR1 has been shown to stimulate keratinocyte migration and facilitate host defense (12).

Low concentrations of COX2 products were observed in the macrophage experiments. Though moderate concentrations of PGD_2_ and PGE_2_ were observed in the HRO/PEHeRo experiments as well as in the stimulated controls, RT-qPCR analyses showed that genes for COX2 were expressed, indicating that the necessary enzymatic machinery for intracellular lipid mediator production was activated. The moderate concentrations of PGD_2_ and PGE_2_ may therefore be attributed to several other factors in the experiment design. Using serum-free media greatly reduces the pool of available arachidonic acid for COX2 conversion as it does not contain lipids. This, combined with moderate endogenous arachidonic acid content in the macrophages (Supplementary Table S1), could have led to a substrate deficiency. Such a potential substrate deficiency in the stimulated control experiments is also supported by the HRO experiments, where a dose-dependent trend for PGD_2_ and PGE_2_ excretion was observed when supplementing the cells with a pool of accessible fatty acids, including arachidonic acid. Furthermore, when M-CSF is used for macrophage differentiation it has previously been reported to yield more M2-like macrophages than when GM-CSF is used, resulting in lower COX2 induction capacity and thus lower concentrations of prostaglandins upon stimulation with LPS (81). Lastly, the LPS concentrations used in the co-stimulation experiments at 1 ng/mL may have been below the threshold for significant COX2 induction (82), as macrophages require higher concentrations of stimuli than undifferentiated monocytes (83,84). This would however not have been the case in the LPS+IFNγ experiments, where the addition of LPS was a 1000-fold higher at 1 µg/mL.

In addition to promoting lipid mediator biosynthesis, we also observed that HRO significantly increased the levels of EPA/DHA in the total fatty acids in the MDMs and significantly increased the levels of EPA/DHA and reduced the levels of AA in the PL-fraction of the fibroblast/keratinocyte co-culture. Allowing for potential replacement of AA in cell membranes could positively impact psoriasis, as overexpressed PLA_2_ enzymes liberating AA for conversion into pro-inflammatory lipid mediators is a driver for psoriatic inflammation (85).

The results presented in this work also show that both HRO and PEHeRo contain SPMs, including: 1) PDX that has been shown to ameliorate inflammation and promote macrophage polarization in experimental models of infection (86). 2) MaR1 that has been reported to display immunoresolving effects in models of skin inflammation through regulation of IL-23 (87), and 3) RvE4 that increases efferocytosis of apoptotic cells (88). The impact of these existing SPMs in the oils was not elucidated in this work, but considering their reported effects, they may provide anti-inflammatory effects and could potentially impact SPM biosynthesis. The effect of exogenous administration of lipid mediators has been shown in humans, where supplementation has been shown to increase blood lipid mediator levels (89). However, as stated above, the treatment of MDMs indicated that the existing lipid mediator levels in HRO and PEHeRo were not the primary drivers for biosynthesis, as the PEHeRo contained far lower quantities of lipid mediators than HRO but resulted in significantly increased levels of DHA-derived SPMs in the cell media.

Four separate findings support that the lipid mediators originated from *de novo* cellular biosynthesis post HRO/PEHeRo stimulation: Firstly, higher quantities of SPMs from the DHA and EPA metabolomes were observed when using PEHeRo compared to HRO. Since PEHeRo contains only 4% - 7% of the monohydroxylated PUFAs present in HRO normalizing lipid mediator output based on the oil content would further amplify the relative difference between the results for HRO and PEHeRo. It is also unlikely that the observed differences in quantities arose from increased autoxidation in PEHeRO compared to HRO, as phospholipid oils in PEHeRo have been reported to be more resilient towards oxidation than oils based on other lipid classes such as the TAGs present in HRO (90,91). Secondly, qualitative profiles also support *de novo* cellular synthesis of the detected lipid mediators: While the lipid mediator profiles of HRO and PEHeRo were similar, they both differed substantially from the profiles observed in the macrophage supernatants. The specific SPMs detected in the macrophage experiments were also inconsistent with autoxidation: trihydroxylated RvD1 and RvD2 were more abundant than the dihydroxylated RvD5, whereas autoxidation reportedly yields higher levels of species with fewer hydroxy groups (92). Thirdly, the PCTRs observed in the experiments were absent from the oils and cannot be formed through autoxidation alone due to their sulfido-conjugated nature. Lastly, genes encoding enzymes required for lipid mediator synthesis were found to be expressed under the experimental conditions. The lack of detection of ALOX15 expression under these conditions was in alignment with literature stating its dependence on IL-4 and IL-13 for expression. However, ALOX15B was observed to be expressed in the macrophages, aligned with what has been previously reported in the absence of IL-4 or IL-13 stimulation (93).

The results from the present study have relevance for understanding the underlying mechanisms for regulation of cell processes involved in the pathophysiology of psoriasis through lipid mediators and SPMs. The key limitation to the data, however, is this being *in vitro* experiments, where all fatty acids and lipid mediators were made very accessible in concentrations that may not reflect realistic *in vivo* concentrations. Thus, the effects of directly metabolically available PUFAs and, particularly, lipid mediators present in the oils may therefore be over-emphasized. It is therefore important to take this into further *in vivo* trials to confirm the relevance of the findings.

The data on skin cells mimicking psoriasis indicates a possible mechanism of resolution of inflammation in the local skin niche which could provide a treatment modality for psoriasis, and it is plausible that these mechanisms also could contribute to resolution of inflammation in other organs and IMIDs. The findings will also contribute to the broader understanding of the pharmacological factors that are in play when administering exogenous sources of SPMs and SPM precursors into biological systems.

In summary, we observed that HRO promotes broad SPM and lipid mediator biosynthesis in macrophages and keratinocytes/fibroblasts, with a subset of SPMs that are consistently upregulated throughout the stimulation/treatment conditions and cell types. Thus, indicating that these pathways may carry significance for signaling and inflammation resolution across several cell types. When treating macrophages with HRO and PEHeRo we found that the macrophages exhibited a dose-dependent trend for biosynthesis of SPMs associated with resolution of inflammation and a shift towards a more protective and possibly reparative phenotype. The presented results on activation of anti-inflammatory lipid mediator pathways are compatible with the beneficial effects on psoriasis observed in clinical trials with HRO and PEHeRo.

### Data availability

Lipid mediator raw data to be shared upon request. Contact corresponding author Thomas Ringheim-Bakka (Arctic Bioscience AS, Industrivegen 42, 6155 Ørsta, Norway) for access.

## Acknowledgments

We would like to thank Daniele Mancinelli and Federico Petrucelli for contributing to providing HRO and PEHeRo and describing the scope of the project, Sissel B. Rønning for contributing to the cell co-culture experiments and scientific discussions, Elin Merete Wetterhus for the skillful technical assistance in analyzing the fatty acid composition of the cells, and Bodil Stige at “Medisinsk biokjemi og blodbank Ålesund” for the provision of buffy coats.

## Abbreviations

AA: arachidonic acid
AcOH: acetic acid
AIM-V: AIM-V medium
ALOX12: arachidonate 12-lipoxygenase
ALOX15: arachidonate 15-lipoxygenase
ALOX15B: arachidonate 15-lipoxygenase type B
ALOX5: arachidonate 5-lipoxygenase
ANOVA: analysis of variance
B2M: beta-2 microglobulin
C18: octadecyl silica gel
CCL2: chemokine ligand 2
cDNA: complementary DNA
COX2: cyclooxygenase-2
CTS: Cell Therapy Systems
CYP450: cytochrome P450
DHA: docosahexaenoic acid
DM: dry matter
DMEM: Dulbeccós modified Eaglés medium
DPA: docosapentaenoic acid
EF1A: eukaryotic translation elongation factor 1 alpha
EPA: eicosapentaenoic acid
Et2O: diethyl ether
EtOH: ethanol
FA: fatty acid
FBS: fetal bovine serum
FID: flame ionization detector
GAPDH: glyceraldehyde-3-phosphate dehydrogenase
GC: gas chromatography
GC-FID: gas chromatography - flame ionization detector
GPCR: G protein-coupled receptor
GPX4: glutathione peroxidase 4
HDHA: hydroxydocosahexaenoic acid
HDPA: hydroxydocosapentaenoic acid
HEPE: hydroxyeicosapentaenoic acid
HETE: hydroxyeicosatetraenoic acid
HPLC: high-performance liquid chromatography
HPRT1: hypoxanthine phosphoribosyltransferase 1
HRO: herring roe oil
HSD: honestly significant difference
ICSR: Immune Cell Serum Replacement
IFN γ: interferon gamma
IFN- γ R1: interferon gamma receptor 1
IHR: immature herring roe
IL: interleukin
IL-17: interleukin 17
IL-17A: interleukin 17A
IL-23: interleukin 23
IMID: immune-mediated inflammatory disease
IS: internal standard
LC-MS/MS: liquid chromatography-tandem mass spectrometry
LC-PUFA: long-chain polyunsaturated fatty acid
LLOQ: lower limit of quantification
LPS: lipopolysaccharide
LTB_4_: leukotriene B_4_
LTC_4_: leukotriene C_4_
LTD_4_: leukotriene D_4_
LTE_4_: leukotriene E_4_
LXA_4_: lipoxin A_4_
MaR1: maresin 1
MaR2: maresin 2
M-CSF: macrophage colony-stimulating factor
MDM: monocyte-derived macrophages
MeOH: methanol
mRNA: messenger RNA
MRM: multiple reaction monitoring
NFKBIZ: NF- κB inhibitor zeta
NMR: nuclear magnetic resonance spectroscopy
PASI: psoriasis area severity index
PBMC: peripheral blood mononuclear cells
PC: phosphatidylcholine
PCR: polymerase chain reaction
PCTR: protectin conjugate in tissue regeneration
PCTR1: protectin conjugate in tissue regeneration 1
PCTR2: protectin conjugate in tissue regeneration 2
PCTR3: protectin conjugate in tissue regeneration 3
PD1: protectin D1
PD1_n-3 DPA_: n-3 docosapentaenoic acid-derived protectin D1
PDX: protectin DX
PEHeRo: phospholipid esters from herring roe
Pen-Strep: penicillin and streptomycin
Ph. Eur.: european pharmacopoeia
PGE3: prostaglandin E3
PLA_2_: phospholipase A_2_
PL: phospholipid
PLS-DA: partial least-squares discriminant analysis
PMN: polymorphonuclear leukocytes
PUFA: polyunsaturated fatty acid
qPCR: quantitative PCR
QTrap: quadrupole ion trap
RBC: red blood cell
REK: regional ethics committee
RNA: ribonucleic acid
RPOL2: RNA polymerase 2
RvD1: resolvin D1
RvD2: resolvin D2
RvD3: resolvin D3
RvD4: resolvin D4
RvD5: resolvin D5
RvD5_n-3 DPA_: n-3 docosapentaenoic acid-derived resolvin D5
RvD6: resolvin D6
RvE1: resolvin E1
RvE2: resolvin E2
RvE3: resolvin E3
RvE4: resolvin E4
RvT1: resolvin T1
RvT2: resolvin T2
RvT4: resolvin T4
S100A7: psoriasine
SEM: standard error of the mean
SPE: solid-phase extraction
SPM: specialized pro-resolving mediators
TAG: triacylglycerol
TLC: thin layer chromatography
UV: ultraviolet
VIP: variable importance in projection
17R-PD1: 17(R)-PD1
17S-HpDHA: 17S-hydroperoxy-DHA)

