## Supplementary material for "Herring roe PLs promote SPM biosynthesis in macrophages and a keratinocyte/fibroblast co-culture as model of psoriasis"

Table S1: Lipid mediator levels (pg/22 mg oil) in HRO and PEHeRo used in the *in vitro* analyses, and % amount of each lipid mediator in PEHeRo compared to in HRO. Data (n=3) is reported as means ± SEM, where “-“ = all replicates below LLOQ.

|  | HRO | | PEHeRo | | |
| --- | --- | --- | --- | --- | --- |
| **DHA-derived mediators** | Mean | SEM | Mean | SEM | % of HRO |
| RvD1 | 464 | 115 | 188 | 18 | 41 % |
| 17R-RvD1 | 293 | 34 | 39 | 1 | 13 % |
| RvD2 | 2 453 | 377 | 350 | 49 | 14 % |
| RvD3 | 3 131 | 556 | - | - | - |
| 17R-RvD3 | 1 353 | 269 | 440 | 23 | 33 % |
| RvD4 | 3 702 | 918 | 615 | 38 | 17 % |
| RvD5 | 86 113 | 6 154 | 6 436 | 203 | 7 % |
| RvD6 | 37 267 | 3 147 | 3 163 | 177 | 8 % |
| PD1 | - | - | - | - | - |
| PDX | 269 899 | 29 684 | 12 134 | 889 | 4 % |
| 17R-PD1 | 32 871 | 1 721 | 2 846 | 126 | 9 % |
| 22-OH-PD1 | - | - | - | - | - |
| PCTR1 | - | - | - | - | - |
| PCTR2 | - | - | - | - | - |
| PCTR3 | - | - | - | - | - |
| MaR1 | 21 771 | 2 047 | 3 679 | 209 | 17 % |
| MaR2 | 17 294 | 1 911 | 1 074 | 29 | 6 % |
| MCTR1 | - | - | - | - | - |
| MCTR2 | - | - | - | - | - |
| MCTR3 | - | - | - | - | - |
| Sum DHA SPMs | 465 213 | 40 227 | 29 333 | 1 479 | 6 % |
| 17-HDHA | 8 004 369 | 801 414 | 459 032 | 16 206 | 6 % |
| 14-HDHA | 3 948 917 | 362 446 | 252 329 | 6 299 | 6 % |
| 13-HDHA | 2 323 637 | 231 368 | 164 919 | 6 366 | 7 % |
| 7-HDHA | 2 370 906 | 312 993 | 158 451 | 4 748 | 7 % |
| 4-HDHA | 6 272 831 | 702 008 | 677 787 | 24 020 | 11 % |
| Sum HDHAs | 22 920 660 | 2 395 069 | 1 712 517 | 56 980 | 7 % |
| DHA | 104 | 4 | 25 | 2 | 24 % |
| **n-3 DPA-derived mediators** |  |  |  |  |  |
| RvT1 | 159 | 25 | 71 | 2 | 45 % |
| RvT2 | - | - | - | - | - |
| RvT3 | - | - | - | - | - |
| RvT4 | 5 855 | 498 | 813 | 41 | 14 % |
| RvD1_n-3 DPA_ | - | - | - | - | - |
| RvD2_n-3 DPA_ | - | - | - | - | - |
| RvD5_n-3 DPA_ | 4 637 | 387 | 379 | 18 | 8 % |
| PD1_n-3 DPA_ | - | - | - | - | - |
| MaR1_n-3 DPA_ | - | - | - | - | - |
| Sum DPA SPMs | 10 651 | 788 | 1 263 | 58 | 12 % |
| 17-HDPA | 993 000 | 43 715 | 37 000 | 529 | 4 % |
| 14-HDPA | 587 000 | 20 075 | 25 750 | 728 | 4 % |
| 13-HDPA | 86 900 | 8 173 | 3 160 | 78 | 4 % |
| 7-HDPA | 4 703 | 929 | 390 | 50 | 8 % |
| Sum HDPAs | 1 671 603 | 72 531 | 66 300 | 1 071 | 4 % |
| DPA | 545 | 14 | 86 | 5 | 16 % |
| **EPA-derived mediators** |  |  |  |  |  |
| RvE1 | - | - | - | - | - |
| RvE2 | - | - | - | - | - |
| RvE3 | - | - | - | - | - |
| RvE4 | 28 982 | 2 786 | 4 380 | 122 | 15 % |
| Sum EPA SPMs | 28 982 | 2 786 | 4 380 | 122 | 15 % |
| 18-HEPE | 11 013 786 | 1 270 530 | 672 777 | 20 065 | 6 % |
| 15-HEPE | 7 687 440 | 831 638 | 328 110 | 11 575 | 4 % |
| 12-HEPE | 7 033 455 | 874 812 | 353 163 | 14 185 | 5 % |
| 11-HEPE | 6 836 150 | 1 006 593 | 283 236 | 8 243 | 4 % |
| 5-HEPE | 8 050 363 | 1 374 487 | 692 706 | 21 735 | 9 % |
| Sum HEPEs | 40 621 194 | 5 314 116 | 2 329 992 | 71 064 | 6 % |
| EPA | 114 298 | 8 831 | 29 739 | 1 801 | 26 % |
| **AA-derived mediators** |  |  |  |  |  |
| LXA_4_ | - | - | - | - | - |
| LXB_4_ | - | - | - | - | - |
| 5S,15S-diHETE | 1 191 | 158 | 187 | 6 | 16 % |
| 15-epi-LXA_4_ | - | - | - | - | - |
| 15-epi-LXB_4_ | - | - | - | - | - |
| LTB4 | 1 951 | 153 | 329 | 9 | 17 % |
| 20-OH-LTB4 | - | - | - | - | - |
| LTC_4_ | - | - | - | - | - |
| LTD_4_ | - | - | - | - | - |
| LTE_4_ | 129 | 59 | 25 | 2 | 19 % |
| Sum | 3 229 | 275 | 541 | 17 | 17 % |
| PGD_2_ | 3 971 | 1 451 | 903 | 56 | 23 % |
| PGE_2_ | 8 678 | 3 367 | 2 098 | 82 | 24 % |
| PGF_2a_ | 359 | 157 | 192 | 8 | 54 % |
| TxB_2_ | 425 | 137 | 129 | 3 | 30 % |
| 20-HETE | 563 320 | 100 475 | 19 673 | 1 861 | 3 % |
| 15-HETE | 147 066 | 15 336 | 6 210 | 288 | 4 % |
| 12-HETE | 61 691 | 3 724 | 3 804 | 161 | 6 % |
| 11-HETE | 34 333 | 4 398 | 2 111 | 86 | 6 % |
| 5-HETE | 531 754 | 51 011 | 34 128 | 1 622 | 6 % |
| Sum HETEs | 1 338 164 | 172 963 | 65 926 | 3 497 | 5 % |
| AA | 121 207 | 8 284 | 20 756 | 1 123 | 17 % |

Table S2: Fatty acid composition of the macrophage cultures shown as % of total fatty acids (FAs). Data (n=3) is reported as means ± SEM, where “-“ = all replicates below LLOQ. *) p < 0.05 (One-factor ANOVA, Tukeys HSD).

|  | Control (% of total FAs) | | | HRO (% of total FAs) | | |
| --- | --- | --- | --- | --- | --- | --- |
|  | Mean |  | SEM | Mean |  | SEM |
| C6:0 | - | ± | - | 0,5 | ± | 0,5 |
| C12:0 | - | ± | - | 0,1 | ± | 0,1 |
| C14:0 | 2,7 | ± | 0,1 | 2,5 | ± | 0,0 |
| C15:0 | 0,2 | ± | 0,0 | 0,3 | ± | 0,0 |
| C16:0 | 23,0 | ± | 0,8 | 21,3 | ± | 1,1 |
| C17:0 | 0,3 | ± | 0,0 | 0,3 | ± | 0,0 |
| C18:0 | 9,5 | ± | 0,8 | 7,6 | ± | 0,5 |
| C24:0 | 2,0 | ± | 0,6 | 1,4 | ± | 0,2 |
| **Sum SFA** | **37,6** | **±** | **2,1** | **36,0** | **±** | **2,2** |
| C15:1 | 0,3 | ± | 0,0 | 0,2 | ± | 0,0 |
| C16:1 | 3,9 | ± | 0,5 | 4,1 | ± | 0,3 |
| C18:1n9 | 26,1 | ± | 0,5 | 23,6 | ± | 2,1 |
| C18:1n7 | 6,9 | ± | 0,9 | 5,9 | ± | 0,2 |
| C20:1n9 | 0,7 | ± | 0,1 | 0,7 | ± | 0,1 |
| C22: n-11 | - | ± | - | 0,2 | ± | 0,0 |
| C22:1 n-9 | 0,3 | ± | 0,1 | 0,3 | ± | 0,0 |
| C24:1 | 0,7 | ± | 0,1 | 0,7 | ± | 0,1 |
| **Sum MUFA** | **38,8** | **±** | **0,3** | **33,8** | **±** | **2,0** |
| C18:2 n-6 | 11,3 | ± | 1,3 | 11,2 | ± | 1,6 |
| C20:2 n-6 | 1,1 | ± | 0,2 | 0,9 | ± | 0,2 |
| C20:3 n-6 | 1,2 | ± | 0,1 | 1,3 | ± | 0,0 |
| C20:4 n-6 | 6,0 | ± | 0,6 | 5,0 | ± | 0,3 |
| **Sum n-6** | **19,6** | **±** | **0,8** | **16,5** | **±** | **1,8** |
| C20:5 n-3 | 0,4 | ± | 0,2 | 1,6^*^ | ± | 0,2 |
| C22:5 n-3 | 1,6 | ± | 0,3 | 3,4^*^ | ± | 0,4 |
| C22:6 n-3 | 1,9 | ± | 1,1 | 6,9^*^ | ± | 1,2 |
| **Sum n-3** | **3,9** | **±** | **1,6** | **13,7^*^** | **±** | **1,7** |
| C22:3/22:4 | 3,0 | ± | 0,0 | 1,1 | ± | 0,4 |

Table S3: Fatty acid composition in the phospholipids of the fibroblast/keratinocyte co-culture stimulated with IL-17A and cultured without (Control) or supplemented with HRO in the growth medium (HRO). Data (n=9) is reported as means ± SEM, where “-” = all replicates below LLOQ. *) p < 0.05 (One-factor ANOVA, Tukeys HSD). 1) Includes C15:0, C17:0, and C22:0.

|  | Control (% of total FAs) | | | HRO (% of total FAs) | | |
| --- | --- | --- | --- | --- | --- | --- |
|  | Mean |  | SEM | Mean |  | SEM |
| C14:0 | 1,4 | ± | 0,0 | 1,2^*^ | ± | 0,0 |
| C15:0 | 0,2 | ± | 0,0 | 0,2 | ± | 0,0 |
| C16:0 | 16,5 | ± | 0,1 | 16,8 | ± | 0,2 |
| C17:0 | 0,2 | ± | 0,0 | 0,3 | ± | 0,0 |
| C18:0 | 8,2 | ± | 0,1 | 10,3^*^ | ± | 0,1 |
| C22:0 | - | ± | - | 0,1 | ± | 0,0 |
| **Sum SFA** | **26,6** | **±** | **0,1** | **28,9^*^** | **±** | **0,3** |
| C14:1 n-5 | - | ± | - | - | ± | - |
| C15:1 | 2,8 | ± | 0,1 | 2,7 | ± | 0,1 |
| C16:1 n-9 | 0,8 | ± | 0,0 | 0,7 | ± | 0,0 |
| C16:1 n-7 | 6,5 | ± | 0,1 | 4,9^*^ | ± | 0,1 |
| C16:1 n-5 | 0,1 | ± | 0,0 | 0,3 | ± | 0,1 |
| C17:1 n-7 | 2,5 | ± | 0,1 | 2,4 | ± | 0,0 |
| C18:1 n-11 | 0,7 | ± | 0,0 | 0,6 | ± | 0,0 |
| C18:1 n-9 | 23,1 | ± | 0,1 | 22,3^*^ | ± | 0,1 |
| C18:1 n-7 | 10,4 | ± | 0,2 | 7,3^*^ | ± | 0,1 |
| C20:1 n-9 | 0,5 | ± | 0,0 | 0,4^*^ | ± | 0,0 |
| C20:1 n-7 | 0,8 | ± | 0,0 | 0,4^*^ | ± | 0,0 |
| C22:1 n-9 | 0,1 | ± | 0,0 | - | ± | - |
| C24:1 n-9 | 0,0 | ± | 0,0 | - | ± | - |
| **Sum MUFA** | **48,3** | **±** | **0,3** | **42,1^*^** | **±** | **0,3** |
| C16:2 n-6 | 0,1 | ± | 0,1 | - | ± | - |
| C18:2 n-6 | 1,5 | ± | 0,0 | 1,5 | ± | 0,0 |
| C18:3 n-6 | 1,2 | ± | 0,0 | 1,1^*^ | ± | 0,0 |
| C20:2 n-6 | 0,2 | ± | 0,1 | -^*^ | ± | - |
| C20:4 n-6 | 3,9 | ± | 0,0 | 3,6^*^ | ± | 0,1 |
| C22:4 n-6 | 2,4 | ± | 0,2 | 2,2 | ± | 0,1 |
| C22:5 n-6 | 0,6 | ± | 0,1 | 0,5 | ± | 0,1 |
| C20:3 n-6 | 0,7 | ± | 0,0 | 0,6 | ± | 0,0 |
| **Sum n-6** | **10,0** | **±** | **0,3** | **8,9^*^** | **±** | **0,1** |
| C16:3 n-4 | 0,3 | ± | 0,1 | 0,4 | ± | 0,1 |
| C16:2 n-3 | 0,4 | ± | 0,1 | 0,4 | ± | 0,1 |
| C20:4 n-3 | 0,6 | ± | 0,1 | 0,4^*^ | ± | 0,1 |
| C20:3 n-3 | 0,1 | ± | 0,1 | - | ± | - |
| C20:5 n-3 | - | ± | - | 0,9^*^ | ± | 0,0 |
| C22:5 n-3 | 1,2 | ± | 0,1 | 2,2 | ± | 0,0 |
| C22:6 n-3 | 1,4 | ± | 0,0 | 3,8^*^ | ± | 0,1 |
| **Sum n-3** | **3,7** | **±** | **0,2** | **7,6^*^** | **±** | **0,1** |

Table S4: Lipid mediator levels (mean pg/3 mL of 4 donors/experiments ± SEM) in supernatants from macrophages incubated with 0.05% HRO prior to stimulation with LPS+IFN-γ. “-“ = all replicates below LLOQ. #) VIP-score >1 in PLS-DA analysis.

|  | Media + LPS + IFN-γ  (Stimulated control) | | HRO 0.05% + LPS + IFN-γ | |
| --- | --- | --- | --- | --- |
| **DHA-derived mediators** | Mean | SEM | Mean | SEM |
| RvD1^#^ | 4,3 | 4,3 | 183,2 | 62,6 |
| 17R-RvD1 | - | - | - | - |
| RvD2 | - | - | 379,6 | 36,6 |
| RvD3 | - | - | 13,0 | 13,0 |
| RvD4^#^ | 13,2 | 8,0 | - | - |
| RvD5 | 1,1 | 1,1 | 11,2 | 6,8 |
| PD1^#^ | - | - | 65,0 | 23,5 |
| PDX^#^ | 22,3 | 10,7 | 215,4 | 27,4 |
| 17R-PD1 | - | - | - | - |
| PCTR1 | - | - | 120,2 | 70,7 |
| PCTR2 | - | - | 191,2 | 116,2 |
| PCTR3 | - | - | - | - |
| MaR1 | - | - | - | - |
| MaR2 | - | - | - | - |
| MCTR1 | - | - | - | - |
| MCTR2 | - | - | 0,8 | 0,8 |
| MCTR3 | - | - | - | - |
| Sum DHA SPMs | 40,9 | 11,2 | 1179,6 | 282,2 |
| **n-3 DPA-derived mediators** |  |  |  |  |
| RvT1 | - | - | - | - |
| RvT2 | - | - | - | - |
| RvT3 | 44,4 | 29,0 | 294,3 | 294,3 |
| RvT4 | - | - | - | - |
| RvD1_n-3 DPA_ | 4,8 | 2,8 | 92,3 | 9,8 |
| RvD2_n-3 DPA_ | - | - | - | - |
| RvD5_n-3 DPA_^#^ | 3,6 | 1,3 | 32,7 | 11,3 |
| PD1_n-3 DPA_^#^ | 106,8 | 68,6 | 875,7 | 318,3 |
| Sum n-3 DPA SPMs | 159,6 | 93,9 | 1295,0 | 466,8 |
| **EPA-derived mediators** |  |  |  |  |
| RvE1 | - | - | - | - |
| RvE2 | 27,1 | 12,4 | 392,0 | 34,8 |
| RvE3 | 2244,8 | 1087,4 | 27937,5 | 1410,5 |
| RvE4 | - | - | 14,3 | 9,0 |
| Sum EPA SPMs | 2271,9 | 1098,9 | 28343,8 | 1393,5 |
| **AA-derived mediators** |  |  |  |  |
| LXA_4_ | - | - | - | - |
| LXB_4_ | - | - | - | - |
| 15-epi-LXA_4_ | 18,3 | 7,4 | 126,6 | 18,9 |
| LTB4 | 9,7 | 5,4 | 75,4 | 39,7 |
| 20-OH-LTB4 | - | - | - | - |
| LTC_4_ | - | - | - | - |
| LTD_4_ | - | - | - | - |
| LTE_4_ | - | - | - | - |
| Sum AA SPMs | 28,0 | 9,6 | 202,0 | 54,8 |
| PGD_2_^#^ | 77,2 | 19,6 | 444,3 | 33,1 |
| PGE_2_^#^ | 2,1 | 2,1 | 35,4 | 3,5 |
| PGF_2a_ | 85,2 | 30,7 | 45,8 | 8,3 |
| TxB_2_ | 4,8 | 4,8 | - | - |

Table S5: SPM levels (mean pg/3 mL of 4 donors/experiments ± SEM) in supernatants from macrophages incubated with HRO/PEHeRo + LPS. “-“ = all replicates below LLOQ. #) VIP-score >1 in PLS-DA analysis.

|  | Media + LPS (Stimulated control) | | | HRO 0.05% + LPS | | HRO 0.25% + LPS | | PEHeRo 0.25% + LPS | |
| --- | --- | --- | --- | --- | --- | --- | --- | --- | --- |
| **DHA-derived mediators** | Mean | | SEM | Mean | SEM | Mean | SEM | Mean | SEM |
| RvD1^#^ | - | | - | - | - | 1401,0 | 469,1 | 3865,8 | 411,0 |
| 17R-RvD1 | - | | - | - | - | - | - | - | - |
| RvD2^#^ | 1,7 | | 1,7 | 2374,8 | 660,7 | 3451,5 | 467,2 | 9307,5 | 965,5 |
| RvD3 | 0,9 | | 0,9 | 72,5 | 72,5 | 435,4 | 150,7 | 390,8 | 390,8 |
| RvD4 | 3,4 | | 3,4 | 2678,8 | 1585,8 | 2328,0 | 2328,0 | 4905,8 | 2962,6 |
| RvD5 | 0,0 | | 0,0 | 317,0 | 25,9 | 573,5 | 87,9 | 27,4 | 27,4 |
| PD1 | 6,1 | | 3,6 | 1183,3 | 418,5 | - | - | - | - |
| PDX | 4,3 | | 2,5 | 1977,3 | 204,3 | 3111,5 | 399,1 | 395,1 | 147,0 |
| 17R-PD1 | - | | - | - | - | - | - | - | - |
| PCTR1 | - | | - | 3,6 | 3,6 | 481,7 | 477,1 | 1771,0 | 1037,2 |
| PCTR2^#^ | 1,7 | | 1,7 | 936,5 | 245,0 | 2004,7 | 723,5 | 5664,0 | 2208,3 |
| PCTR3 | - | | - | 24,0 | 17,8 | 96,1 | 54,4 | 6,1 | 6,1 |
| MaR1 | - | | - | - | - | - | - | - | - |
| MaR2 | - | | - | 43,2 | 43,2 | 74,2 | 74,2 | - | - |
| MCTR1 | - | | - | - | - | - | - | - | - |
| MCTR2 | - | | - | 1,7 | 1,7 | 18,8 | 11,8 | - | - |
| MCTR3 | - | | - | 97,2 | 71,5 | 418,3 | 143,4 | - | - |
| Sum DHA SPMs | 18,1 | | 6,3 | 9709,7 | 1242,7 | 14394,6 | 2921,7 | 26333,4 | 2893,4 |
| **n-3 DPA-derived mediators** |  | |  |  |  |  |  |  |  |
| RvT1^#^ | - | | - | 10,6 | 1,9 | 24,8 | 2,5 | 9,3 | 5,6 |
| RvT2^#^ | - | | - | 27,0 | 27,0 | 49,7 | 49,7 | 281,2 | 22,7 |
| RvT3 | - | | - | 1080,0 | 1080,0 | - | - | 7332,5 | 4330,6 |
| RvT4 | - | | - | 14,9 | 14,9 | 105,0 | 18,2 | 9,6 | 9,6 |
| RvD1_n-3 DPA_^#^ | 1,0 | | 0,3 | 197,6 | 72,8 | 315,6 | 116,3 | 523,6 | 108,1 |
| RvD2_n-3 DPA_ | - | | - | - | - | - | - | - | - |
| RvD5_n-3 DPA_ | 0,4 | | 0,3 | 156,2 | 36,5 | 214,5 | 38,5 | 54,8 | 5,0 |
| PD1_n-3 DPA_ | - | | - | 2575,0 | 408,8 | 1688,8 | 1163,3 | 655,5 | 420,5 |
| Sum DPA SPMs | 1,4 | | 0,5 | 4061,3 | 1474,4 | 2398,4 | 1265,7 | 8866,6 | 4525,5 |
| **EPA-derived mediators** |  | |  |  |  |  |  |  |  |
| RvE1 | - | | - | - | - | - | - | - | - |
| RvE2^#^ | 2,4 | 0,5 | | 3045,0 | 381,6 | 6860,5 | 402,7 | 4091,3 | 391,3 |
| RvE3^#^ | 42,2 | 24,4 | | 177550,0 | 39519,4 | 269325,0 | 24428,0 | 396475,0 | 73593,3 |
| RvE4 | - | - | | 48,6 | 48,6 | 194,6 | 121,6 | 87,8 | 50,9 |
| Sum EPA SPMs | 44,6 | 24,1 | | 180643,6 | 39882,0 | 276380,1 | 24166,6 | 400654,0 | 73316,3 |
| **AA-derived mediators** |  |  | |  |  |  |  |  |  |
| LXA_4_ | - | - | | - | - | - | - | - | - |
| LXB_4_ | - | - | | - | - | - | - | - | - |
| 15-epi-LXA_4_ | 8,9 | 3,1 | | - | - | - | - | 379,0 | 225,6 |
| LTB4 | 3,7 | 3,7 | | 25,3 | 25,3 | 88,0 | 50,8 | 178,8 | 149,7 |
| 20-OH-LTB4 | - | - | | - | - | - | - | - | - |
| Sum AA SPMs | 12,6 | 6,1 | | 25,3 | 25,3 | 88,0 | 50,8 | 557,8 | 304,7 |
| LTC_4_ | - | - | | - | - | - | - | - | - |
| LTD_4_ | - | - | | - | - | - | - | - | - |
| LTE_4_ | - | - | | - | - | - | - | - | - |
| PGD_2_^#^ | 31,9 | 3,6 | | 792,3 | 220,9 | 1277,8 | 112,8 | 1531,3 | 344,9 |
| PGE_2_^#^ | 1,0 | 0,6 | | 232,0 | 9,3 | 493,6 | 83,1 | 229,4 | 88,2 |
| PGF_2a_^#^ | 39,3 | 12,7 | | 48,4 | 6,9 | 80,6 | 11,7 | 38,5 | 13,2 |
| TxB_2_ | - | - | | - | - | - | - | - | - |

Table S6: Sum SPMs and SPM precursor levels (mean of 3 biological replicates ± SEM) in fibroblast/keratinocyte coculture cell supernatants (pg/mL) incubated with IL-17A (control), HRO, HRO+IL-17A, or untreated. SPMs were identified and quantified using LC-MS/MS. “-“ = all replicates below LLOQ.

| **Lipid mediators** | Control | | Media + IL-17A (Stim. Control) | | HRO | | IL-17A + HRO | |
| --- | --- | --- | --- | --- | --- | --- | --- | --- |
| **DHA-derived mediators** | Mean | SEM | Mean | SEM | Mean | SEM | Mean | SEM |
| RvD1 | - | - | - | - | 1,8 | 0,9 | 4,4 | 2,5 |
| 17R-RvD1 | - | - | - | - | - | - | - | - |
| RvD2 | - | - | - | - | 15,8 | 1,6 | 4,3 | 4,3 |
| RvD3 | 6,3 | 6,3 | 11,3 | 11,3 | - | - | 19,8 | 13,0 |
| 17R-RvD3 | - | - | - | - | 1,7 | 1,7 | 2,2 | 1,1 |
| RvD4 | - | - | - | - | - | - | - | - |
| RvD5 | - | - | - | - | 4,1 | 0,4 | 4,0 | 0,5 |
| RvD6 | 0,2 | 0,2 | - | - | 16,5 | 1,1 | 10,3 | 1,4 |
| PD1 | - | - | - | - | 19,4 | 2,6 | 6,7 | 6,7 |
| PDX | - | - | - | - | - | - | - | - |
| 17R-PD1 | - | - | - | - | - | - | - | - |
| PCTR1 | 0,1 | 0,1 | - | - | - | - | 0,1 | 0,1 |
| PCTR2 | - | - | - | - | - | - | - | - |
| PCTR3 | - | - | - | - | - | - | - | - |
| MaR1 | - | - | - | - | - | - | - | - |
| MaR2 | - | - | - | - | 0,3 | 0,3 | - | - |
| MCTR1 | 0,3 | 0,3 | 0,3 | 0,3 | - | - | 1,0 | 0,5 |
| MCTR2 | - | - | - | - | - | - | - | - |
| MCTR3 | - | - | - | - | - | - | - | - |
| Sum | 6,8 | 6,0 | 11,6 | 11,6 | 59,6 | 1,8 | 52,9 | 14,5 |
| **n-3 DPA-derived mediators** |  |  |  |  |  |  |  |  |
| RvT1 | - | - | - | - | - | - | 1,2 | 0,6 |
| RvT2 | - | - | 0,1 | 0,1 | - | - | - | - |
| RvT3 | - | - | - | - | - | - | - | - |
| RvT4 | - | - | - | - | - | - | - | - |
| RvD1_n-3 DPA_ | - | - | - | - | - | - | - | - |
| RvD2_n-3 DPA_ | - | - | - | - | - | - | - | - |
| RvD5_n-3 DPA_ | - | - | - | - | 0,2 | 0,2 | 0,3 | 0,3 |
| PD1_n-3 DPA_ | - | - | - | - | - | - | - | - |
| 22-OH-PD1n-3 DPA | - | - | - | - | - | - | - | - |
| MaR1n-3 DPA | 26,8 | 26,8 | 26,6 | 26,6 | 41,7 | 21,0 | - | - |
| MaR2n-3 DPA | - | - | - | - | - | - | - | - |
| Sum | 26,8 | 26,8 | 26,7 | 26,5 | 41,8 | 21,1 | 1,5 | 0,8 |
| **EPA-derived mediators** |  |  |  |  |  |  |  |  |
| RvE1 | - | - | - | - | - | - | - | - |
| RvE2 | 0,1 | 0,1 | 0,1 | 0,1 | 34,6 | 1,0 | 31,9 | 3,9 |
| RvE3 | - | - | - | - | 1562,3 | 38,9 | 1292,7 | 92,1 |
| RvE4 | 0,1 | 0,1 | - | - | - | - | - | - |
| Sum | 0,2 | 0,1 | 0,1 | 0,1 | 1596,9 | 39,4 | 1324,6 | 95,1 |
| **AA-derived mediators** |  |  |  |  |  |  |  |  |
| LXA_4_ | - | - | - | - | - | - | - | - |
| LXB_4_ | - | - | - | - | - | - | - | - |
| 15-epi-LXA_4_ | - | - | - | - | - | - | - | - |
| 15-epi-LXB4 | - | - | - | - | - | - | - | - |
| 5S,15S-diHETE | - | - | - | - | - | - | - | - |
| LTB4 | 0,1 | 0,1 | 0,2 | 0,1 | 0,4 | 0,2 | - | - |
| 20-OH-LTB4 | - | - | - | - | - | - | - | - |
| 20-COOH-LTB4 | - | - | - | - | - | - | - | - |
| LTC_4_ | - | - | - | - | - | - | - | - |
| LTD_4_ | - | - | - | - | - | - | - | - |
| LTE_4_ | - | - | - | - | - | - | - | - |
| PGD_2_ | 4,9 | 0,5 | 5,9 | 0,2 | 6,6 | 0,3 | 5,7 | 0,5 |
| PGE_2_ | 3,9 | 3,2 | 57,8 | 5,8 | 3,8 | 2,0 | 41,2 | 6,3 |
| PGF_2a_ | 4,7 | 0,2 | 8,8 | 0,5 | 6,0 | 0,1 | 10,1 | 1,2 |
| TxB_2_ | 17,0 | 1,4 | 15,8 | 0,6 | 18,8 | 0,1 | 16,3 | 1,6 |
| Sum | 30,7 | 4,4 | 88,5 | 6,7 | 35,7 | 2,1 | 73,2 | 9,4 |
| **Monohydroxy-DHAs** |  |  |  |  |  |  |  |  |
| 17-HDHA | 9,7 | 0,7 | 9,4 | 0,8 | 157,4 | 3,3 | 154,8 | 13,4 |
| 14-HDHA | 5,2 | 1,0 | 6,0 | 0,6 | 68,6 | 0,7 | 67,0 | 3,0 |
| 13-HDHA | 4,5 | 0,5 | 4,8 | 0,3 | 52,7 | 0,8 | 56,0 | 3,7 |
| 7-HDHA | 3,1 | 0,6 | 2,8 | 0,5 | 52,2 | 2,1 | 50,0 | 1,0 |
| 4-HDHA | 1,4 | 1,4 | 2,7 | 1,4 | 49,9 | 4,0 | 67,0 | 5,0 |
| Sum HDHAs | 24,0 | 4,1 | 25,7 | 3,0 | 380,8 | 5,7 | 394,7 | 20,3 |
| **Monohydroxy-DPAs** |  |  |  |  |  |  |  |  |
| 17-HDPA | 4,1 | 2,1 | 3,9 | 1,9 | 28,5 | 0,4 | 36,5 | 4,3 |
| 14-HDPA | - | - | - | - | - | - | - | - |
| 13-HDPA | - | - | - | - | - | - | - | - |
| 7-HDPA | - | - | 0,4 | 0,4 | 0,9 | 0,2 | 0,3 | 0,3 |
| Sum HDPAs | 4,1 | 2,1 | 4,3 | 1,6 | 29,4 | 0,4 | 36,8 | 4,0 |
| **Monohydroxy-EPAs** |  |  |  |  |  |  |  |  |
| 18-HEPE | 20,3 | 1,0 | 17,2 | 0,5 | 945,1 | 36,7 | 766,8 | 11,5 |
| 15-HEPE | 2,5 | 0,3 | 2,1 | 0,1 | 37,5 | 1,0 | 35,8 | 4,0 |
| 12-HEPE | 2,4 | 0,2 | 1,6 | 0,8 | 51,2 | 1,5 | 43,3 | 2,0 |
| 11-HEPE | 1,1 | 0,2 | 0,9 | 0,1 | 25,7 | 0,6 | 24,4 | 1,9 |
| 5-HEPE | 4,1 | 0,6 | 3,5 | 0,3 | 96,7 | 2,5 | 87,0 | 2,1 |
| Sum HEPEs | 30,4 | 2,0 | 25,4 | 1,0 | 1156,1 | 37,1 | 957,2 | 4,4 |
| **Monohydroxy-AAs** |  |  |  |  |  |  |  |  |
| 20-HETE | 21,4 | 10,9 | 24,2 | 4,7 | 321,6 | 12,1 | 264,4 | 22,2 |
| 15-HETE | 10,4 | 0,9 | 11,2 | 0,3 | 13,0 | 0,7 | 15,3 | 2,6 |
| 12-HETE | 12,1 | 1,1 | 13,0 | 0,4 | 17,6 | 0,4 | 17,9 | 1,7 |
| 11-HETE | 3,8 | 0,2 | 5,2 | 0,2 | 4,7 | 0,2 | 6,3 | 1,1 |
| 5-HETE | 7,4 | 0,3 | 8,5 | 0,2 | 15,1 | 0,8 | 15,8 | 1,0 |
| Sum HETEs | 55,0 | 12,7 | 62,1 | 4,6 | 372,1 | 13,5 | 319,8 | 22,4 |

Table S7: Primers used in expression analyses of enzymes in macrophage experiments.

| COX2, PTGS2 | hs_115_PTGS2_F2 | hs_116_PTGS2_R2 |
| --- | --- | --- |
| ALOX5, 5LOX | hs_369_ALOX5_F | hs_370_ALOX5_R |
| ALOX12, 12LOX | hs_111_ALOX12_F2 | hs_112_ALOX12_R2 |
| ALOX15, 15LOX, | hs_173_15LOX_F1 | hs_174_15LOX_R1 |
| ALOX15B | hs_363_ALOX15B_F | hs_364_ALOX15B_R |

Table S8: Primers used in expression analyses of enzymes in keratinocytes and fibroblasts.

| Gene | Nofima Primer name (can be removed) | 5’-3’ | Genbank |
| --- | --- | --- | --- |
| COX2, PTGS2 | hs_115_PTGS2_F2 | GGCCATGGGGTGGACTTAAA | NM_000963 |
|  | hs_116_PTGS2_R2 | ACCGTAGATGCTCAGGGACT |  |
| ALOX5, 5LOX | hs_369_ALOX5_F | CATGCCCTCCTACACGGTC | NM_000698.5 |
|  | hs_370_ALOX5_R | TATGAATCCACCGCGCCAC |  |
| ALOX12, 12LOX | hs_111_ALOX12_F2 | TCTGGAGATGGCCCTCAAAC | NM_000697 |
|  | hs_112_ALOX12_R2 | GGGTTGGCACCATTGAGGAA |  |
| ALOX15, 15LOX, | hs_173_15LOX_F1 | GCTGGAAGGATCTAGATGACT | M23892.1 |
|  | hs_174_15LOX_R1 | TGGCTACAGAGAATGACGTTG |  |
| ALOX15B | hs_363_ALOX15B_F | CCTGCACATCAACACACTCG | NM_001141.3 |
|  | hs_364_ALOX15B_R | TTCAGAGACAAAGCGTTCCA |  |
| EF1a | hs_31_RPOL2_F2 | GACACGTAGATTCGGGCAAGTCCA | NM_001402.5 |
|  | hs_32_RPOL2_R2 | CCATCTCAGCAGCCTCCTTCTCAA |  |
| RPOL2 | hs_31_RPOL2_F2 | GCGCAATGAGCAGAACGGCG | NM_000937.4 |
|  | hs_32_RPOL2_R2 | ACTTCTGCATGGCACGGGGC |  |
| GAPDH | hs_33_GAPDH_F1 | ATCCCATCACCATCTTCCAGGAGC | NM_002046.3 |
|  | hs_34_GAPDH_R1 | AAATGAGCCCCAGCCTTCTCCAT |  |

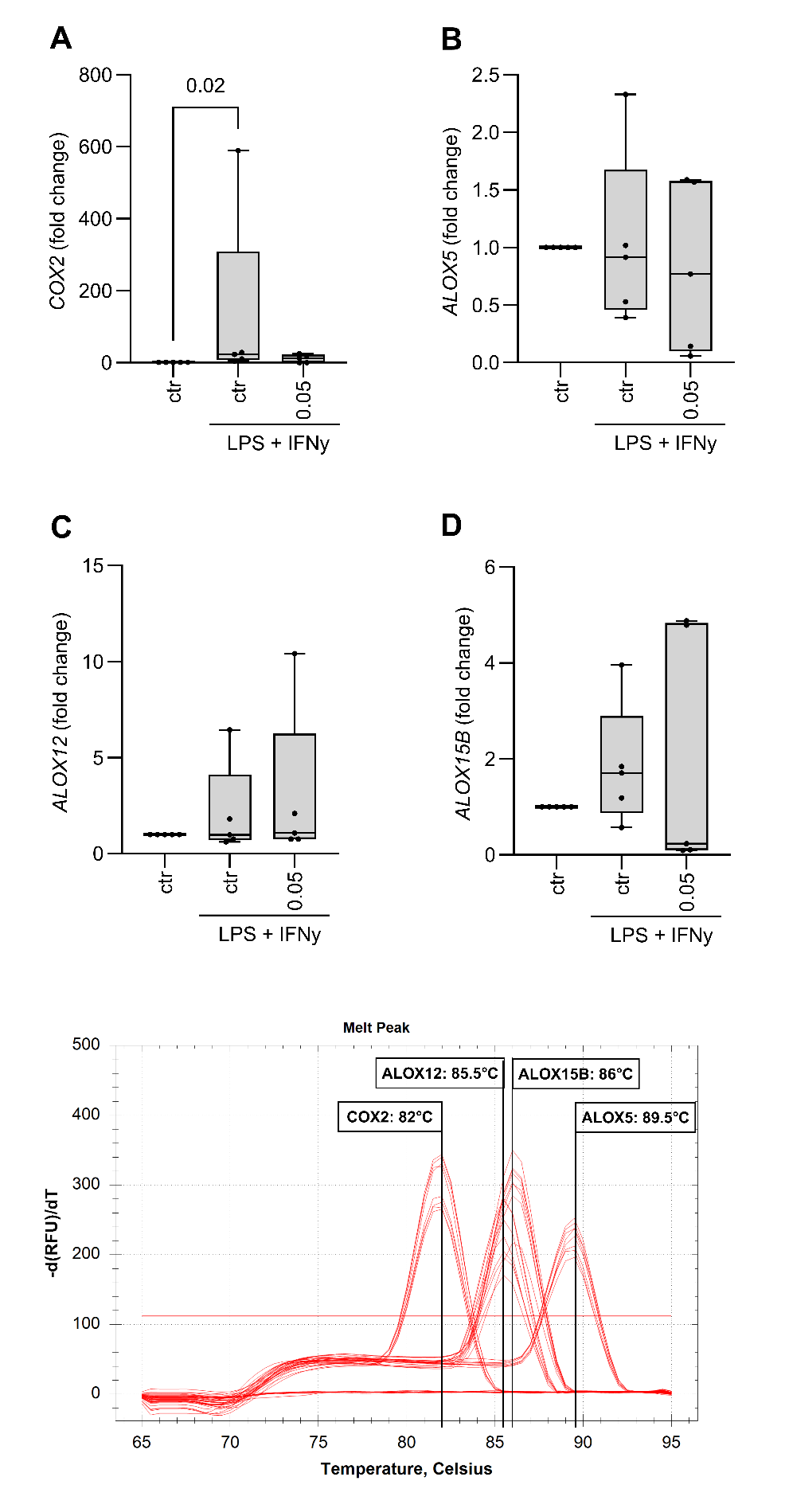
Figure S1: Expression of genes in MDMs for enzymes A) COX2, B) ALOX5, C) ALOX12, and ALOX15B in unstimulated control, stimulated control (LPS + IFNγ), and treated with HRO (0.05%) + LPS + IFNγ.

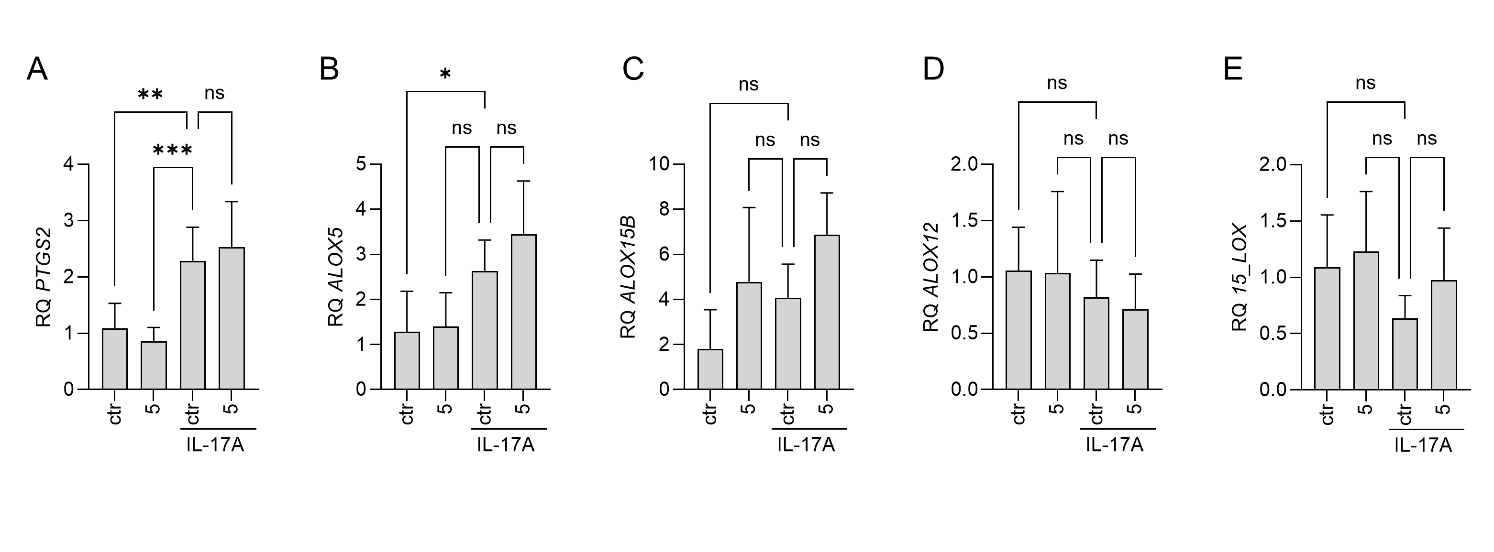
Figure S2: Expression of genes in keratinocyte/fibroblast co-culture for enzymes A) COX2, B) ALOX5, C) ALOX15B, D) ALOX12, and 15LOX in unstimulated control, treated with HRO (5 µg/mL), stimulated control (IL-17A), and treated with HRO (5 µg/mL) + IL-17A. Graphs are generated from data based on 2 technical replicates from 3 biological replicates per group. Graphs show relative gene expression compared to ctr. *) p < 0.05, **) p < 0.01, and ***) p < 0.001 (One-way ANOVA compared to mean of ctr + IL-17A)
